# Role of GM-CSF in regulating metabolism and mitochondrial functions critical to macrophage proliferation

**DOI:** 10.1101/2021.02.10.430444

**Authors:** Matthew Wessendarp, Miki Watanabe, Serena Liu, Traci Stankiewicz, Yan Ma, Kenjiro Shima, Claudia Chalk, Brenna Carey, Lindsey-Romick Rosendale, Marie Dominique Filippi, Paritha Arumugam

## Abstract

Granulocyte-macrophage colony-stimulating factor (GM-CSF) exerts pleiotropic effects on macrophages and is required for self-renewal but the mechanisms responsible are unknown. Using GM-CSF receptor-β-chain deficient (*Csf2rb*^*−/−*^) mice, we show GM-CSF is critical for mitochondrial turnover, functions, and integrity. GM-CSF signaling is essential for fatty acid β-oxidation and markedly increased tricarboxylic acid cycle activity, oxidative phosphorylation, and ATP production. GM-CSF also regulated cytosolic pathways including glycolysis, pentose phosphate pathway, and amino acid synthesis. We conclude that GM-CSF regulates macrophages in part through a critical role in maintaining mitochondria, which are necessary for cellular metabolism as well as proliferation and self-renewal.

## 1.0 Introduction

Alveolar macrophages (AMs) have recently been identified as a self-renewing cell population for which long-term maintenance does not require replenishment from non-pulmonary sources in healthy adult mice (Ginhoux et al., 2016; Hashimoto et al., 2013; Sieweke and Allen, 2013); however, the mechanisms governing AM self-renewal are unknown. Hematopoietic stem cells (HSC) undergo self-renewal and their long-term maintenance has been well studied and demonstrated to depend on mitochondria biogenesis (Basu et al., 2013; Cunningham et al., 2007; Takihara et al., 2019), mitophagy (Ito et al., 2016), mitochondrial metabolism and energy production (Ito et al., 2012; Ito and Ito, 2018; Janzen et al., 2006; Manesia et al., 2015), and asymmetric cell division (Hinge et al., 2020; Katajisto et al., 2015; Takihara et al., 2019). The proliferation of tissue-resident macrophages can be stimulated by a number of molecules including endogenous cytokines (granulocyte/macrophage-colony stimulating factor (GM-CSF), macrophage-colony stimulating factor (M-CSF), interleukin-4 (IL-4), IL-34 (Hashimoto et al., 2013; Pridans et al., 2018; Wang et al., 2012), other biomolecules (neuropeptide FF, nuclear retinoid X receptor, adiponectin, oxidized low-density lipoprotein), *MafB* levels (Aziz et al., 2009; Soucie et al., 2016), and by activation of various normal macrophage functions (Fc-receptor-mediated phagocytosis and complement-mediated activation (Hui et al., 2015; Luo et al., 2005; Menendez-Gutierrez et al., 2015; Senokuchi et al., 2005; Wang et al., 2012; Waqas et al., 2017). Notwithstanding, the precise role of any of these molecules and underlying mechanisms in regulating AM self-renewal, is unclear.

GM-CSF has long been known to stimulate proliferation of human AMs (Nakata et al., 1991) and numerous studies show GM-CSF regulates the survival, proliferation, differentiation, and functions of AMs in mice and humans (Berclaz et al., 2007; Berclaz et al., 2002a; Berclaz et al., 2002b; Bonfield et al., 2003; Carey et al., 2007; Gonzalez-Juarrero et al., 2005; Nakata et al., 1991; Shibata et al., 2001; Suzuki et al., 2014; Uchida et al., 2009; Uchida et al., 2004). Constitutive over-expression of GM-CSF in the lungs of transgenic mice stimulates uncontrolled proliferation and expansion resulting in massive intra-alveolar AM accumulation – a lung phenotype bearing a striking resemblance to a human lung disease, desquamative interstitial pneumonitis (DIP), which is characterized by massive AM accumulation, emphysema, increased morbidity, and increased mortality (Suzuki et al., 2020). Systemic overexpression of GM-CSF in mice results in marked accumulation of macrophages in tissues throughout the body and increased mortality (Lang et al., 1987). In contrast, disruption of GM-CSF signaling results in another lung disease, pulmonary alveolar proteinosis (PAP), characterized by functional impairment of AMs including reduced pulmonary surfactant clearance and numerous other functions (Arumugam et al., 2019; Suzuki et al., 2014; Suzuki et al., 2008; Suzuki et al., 2010; Trapnell and Whitsett, 2002; Willinger et al., 2011). Of note, the manifestations of this lung disease are similar regardless of whether disruption of GM-CSF signaling occurs by ablation of the gene encoding GM-CSF, its receptor alpha or beta subunits, or neutralizing GM-CSF autoantibodies (Arumugam et al., 2019; Berclaz et al., 2007; Dranoff et al., 1994; Shibata et al., 2001; Suzuki et al., 2014; Suzuki et al., 2011; Tanaka et al., 2011). Disruption of GM-CSF signaling in mice, monkeys, and man is associated with decreased expression of the master transcriptional regulator, PU.1, in all three species, as well as the downstream transcription factor, PPARγ that is required for the AM differentiation and specification (Berclaz et al., 2002b; Bonfield et al., 2003; Carey et al., 2007; Gonzalez-Juarrero et al., 2005; Sakagami et al., 2010; Sallese et al., 2017; Suzuki et al., 2014; Suzuki et al., 2011; Suzuki et al., 2008; Suzuki et al., 2010; Trapnell and Whitsett, 2002; Uchida et al., 2009; Uchida et al., 2004). Disruption of GM-CSF signaling is also associated with marked downregulation of the expression of genes required for fatty acid metabolism including carnitine palmitoyltransferase 1 (*CPT-lα*), *CPT-2*, and fatty acid binding protein-1 (*FABP-1*), suggesting GM-CSF may be required for fatty acid metabolism in macrophages (Suzuki et al., 2014).

Transplantation of healthy bone marrow-derived macrophages into the lungs of GM-CSF receptor alpha (*Csf2ra*^*−/−*^) or beta deficient (*Csf2rb*^*−/−*^) mice without myeloablation results in their adoption of a normal AM phenotype, engraftment, replacement of the endogenous (dysfunctional) AMs, and long-term persistence (Suzuki et al., 2014). This model system has been extensively studied in a program to develop this approach – pulmonary macrophage transplantation (PMT) – as therapy of hereditary PAP caused by genetic mutations in *CSF2RA or CSF2RB.* A cornerstone of the treatment approach is that the transplanted macrophages self-renew, thereby contributing to the long-term persistence and durability of the treatment (Arumugam et al., 2019; Suzuki et al., 2014). These studies also identified a reciprocal feedback loop through which pulmonary GM-CSF has been demonstrated to regulate the size of the AM population in mice (Suzuki et al., 2014). Despite its obvious importance to PMT therapy, the mechanism(s) regulating self-renewal and persistence of the transplanted macrophages are unknown.

Based on these findings, we hypothesized that GM-CSF regulates the proliferation of AMs by modulating macrophage metabolism and energy production. Our experimental approach employed genetically modified mice deficient in GM-CSF receptor expression to study cell cycle progression/proliferation assays, nuclear magnetic resonance (NMR)-based metabolomics (Matrka et al., 2017), RNA-sequencing-based transcriptomics, flow cytometry-based marker analysis for determining mitochondrial mass and functions (Hinge et al., 2020), confocal- and electron microscopic-based morphological/ultrastructural analysis, and Seahorse-based measurements of metabolic activity (glycolysis and oxidative phosphorylation) and fatty acid oxidation (FAO) analysis (Fletcher et al., 2015; Huang et al., 2014; Ito et al., 2012). Results show that GM-CSF stimulation is required for the maintenance of mitochondrial mass and function in macrophages, as well as for regulation of glucose and fatty acid catabolism, amino acid and nucleotide precursor synthesis, energy production, and cellular proliferation, all of which are relevant to AM self-renewal.

## 2.0 Materials and methods

### 2.1. Mice

The B6.SJL-*Pfprc*^*a*^*Pepc*^*b*^/BoyJ (Stock No: 002014/B6 CD45.1) and *Csf2rb*^*−/−*^ mice used in this study were bred and experiments conducted in Cincinnati Children’s Hospital Research Foundation (CCHRF) vivarium using protocols approved by the institutional animal care and use committee (IACUC) at Cincinnati Children’s Hospital Medical Center. The generation of *Csf2rb*^*−/−*^ mice has been described previously (Robb et al., 1995).

### 2.2. Preparation of bone marrow derived macrophages (BMDMs)

Mononuclear cells were isolated from the bone marrow of WT or *Csf2rb*^*−/−*^ mice using Histopaque-1083 (Sigma) by centrifugation at room temperature for 30 minutes. The buffy coat comprising of the mononuclear cells were washed with DPBS and resuspended in DMEM containing 10% heat-inactivated fetal bovine serum (FBS), 1X penicillin/streptomycin, 10ng/mL recombinant mouse GM-CSF (R & D systems) and 10ng/mL recombinant mouse M-CSF (R & D systems). The cells were plated in a 100mm dish and incubated in tissue-culture incubator for 2-3 hours. After the initial incubation period, the nonadherent cell fractions were collected from the dishes by gentle rinsing and seeded into a new 100mm dish for expansion and differentiation to BMDMs. The cells were cultured for 7 days with media changes at every 2-3 days. The adherent cells were collected on day 7 using TrypLE select enzyme (Fisher Scientific) with 0.3% pluronic acid. The harvested BMDMs were washed with DPBS and used for experiments.

### 2.3. Collection of alveolar macrophages

Alveolar macrophages were collected from the lung surface of mice by bronchoalveolar lavage (BAL) as described previously (LeVine et al., 1999). The cells were concentrated by centrifugation at 1100 rpm for 10 minutes at room temperature.

### 2.4. Measurement of mitochondrial mass

Alveolar macrophages (50,000 cells) were sedimented onto slides (Cytospin, Shandon Inc) by gentle centrifugation at 400 rpm for 4 minutes at room temperature using a Cytospin 4 Cytocentrifuge. Following this, the macrophages were fixed with 4% paraformaldehyde, washed with DPBS, permeabilized using 0.1% Triton X-100 for 5 minutes at room temperature, rinsed with DPBS, and blocked for 20 min with 2% bovine serum albumin (BSA). After blocking, the slides were incubated with TOMM20 antibody for 2-4 hours at room temperature, washed with PBS and secondary antibodies were added. Fluorescently labeled lectin and wheat germ agglutinin (WGM) were used to stain the cell membrane of alveolar macrophages. Counterstaining was done with DAPI. Images were acquired using confocal microscope and measurement of TOMM20 mean fluorescence intensity was done using NIS elements software.

### 2.5. Cell cycle analysis

Hoechst 33342 stain (ThermoFisher Scientific) was used for cell cycle analysis. Approximately 0.5-1×10^6^ cells were stained with 2μg/mL Hoechst dye and the cells were incubated for 60 minutes at 37°C in complete medium. After incubation, the cells were washed, and the fluorescence was measured using ultra-violet and 488nm excitation in a LSRII flow cytometer (BD Sciences) instrument.

### 2.6. Measurement of Macrophage Proliferation

For the cell proliferation assay, 3×10^4^-5×10^4^ macrophages were plated in a 96 well plate in a 100ul volume of complete cell culture medium. The cell proliferation reagent WST-1 (Sigma, St. Louis) was added to the culture medium (1:10 final dilution) for 4 hours at 37°C. Additional wells were plated with the same volume of culture medium and WST-1 reagent for use as blank wells for background absorbance correction. After the incubation period, the absorbance of the samples was measured against the background control between 440 nm using an ELISA reader (Synergy HTX multimode reader, BioTek, Winooski, VT).

Realtime-Glo MT Cell Viability assay: The assay was set up as a continuous read format according to the manufacturer’s instructions (Promega, Madison, WI) in a 96 well format using 5,000 cells/well in a 100ul volume. The MT cell viability substrate and the NanoLuc enzyme were added to the cell suspension and the cells were incubated in a tissue culture incubator until measurements. The reducing potential of macrophages was assessed by measuring the luminescence at different time points. The reducing potential of the cell was used as a metabolic marker of cell viability.

### 2.7. Cellular and mitochondrial ROS measurement

For analysis of intracellular ROS, macrophages were incubated with 5 μM 2’, 7’ dichlorofluorescein diacetate (DCFDA, Invitrogen at 37°C for 30 min, followed by acquisition using FACS Canto I (BD Sciences) within 30 minutes of staining. The level of fluorescent 2’, 7’ dichlorofluorescein (DCF) detected was used as a measure of intracellular ROS.

Mitochondrial superoxide (Mitosox) measurement was done by incubating macrophages with 5uM MitoSox Red fluorogenic dye specifically targeted to mitochondria in live cells. The cells were stained in dark a 500ul volume for 30 minutes at 37°C, washed, and the mean fluorescence intensity of the oxidized MitoSox red was measured using FACS Canto I within 30 minutes of staining.

### 2.8. Measurement of Mitochondrial Membrane Potential (MMP)

Tetramethyl rhodamine ethyl ester (TMRE) was used to assess the MMP according to manufacturer’s instructions. Briefly, approximately 2×10^5^-5×10^5^ macrophages were stained in a 500ul volume with 200nM TMRE for 15-30 minutes at 37°C and 5% CO_2_ followed by flow cytometry to measure the mitochondrial membrane potential.

### 2.9. Measurement of cellular respiration

The bioenergetic profile of BMDMs were assessed by measuring the oxygen consumption rate (OCR) and extracellular acidification rate (ECAR) using a XF cell Mito Stress Test kit and XF glycolysis stress kit (Agilent Technologies) as per manufacturer’s instructions using a Seahorse XF extracellular flux analyzer (Agilent technologies, North Billerica, Billerica, MA). Briefly, 50,000 BMDMs from WT and *Csf2rb*^*−/−*^ mice were plated in Seahorse XF96 microplates (Seahorse Biosciences, Europe) and incubated for 12 hours in DMEM medium containing 10% fetal bovine serum (FBS), 1% Penicillin/streptomycin solution, 10ng/mL recombinant mouse GM-CSF and 10ng/mL recombinant mouse M-CSF. Prior to assay, the cell culture growth medium is replaced with prewarmed Seahorse XF base medium and the plate is placed into a 37°C, non-CO_2_ incubator for 45-60 minutes. The assay medium for measuring OCR contains 10 mM glucose, 1mM sodium pyruvate, and 2mM glutamine (Agilent Technologies). The first step is to measure the basal respiration in macrophages prior to the addition of Oligomycin A (OM). Sequential addition of inhibitors (Oligomycin A: 1.5μM, carbonyl cyanide 4-(trifluoromethoxy) phenyl-hydrazone (FCCP: 1.0 μM) and Rotenone/Antimycin A (R&AA: 0.5 μM) of the electron transport chain allow for the calculation of OCR-linked ATP production, maximal respiration capacity, and spare respiratory capacity. Extracellular acidification rate (ECAR) allows measurement of lactic acid levels formed during the conversion of glucose to lactate during the anaerobic glycolysis. The following compounds are added in sequence: Glucose (10mM), Oligomycin (1.0 μM) and 2-deoxy glucose (50mM), to enable measurements of glycolysis, glycolytic capacity, and glycolytic reserve measurements, respectively.

### 2.10. Fatty acid oxidation measurement

The ability of the cells to oxidize exogenous fatty acid substrates was measured using the Agilent Seahorse XF Cell Mito Stress Test kit according to manufacturer’s recommendations. BMDMs (4×0^5^ cells) were seeded in Agilent Seahorse XF96 cell culture microplate and allowed to adhere in complete culture medium. The growth medium was then replaced and incubated for 24 hours with substrate limited DMEM medium containing reduced concentrations of glucose (0.5mM), glutamax (1mM) and fetal bovine serum (1%) to deplete endogenous substrates within the cells. Carnitine hydrochloride (0.5mM) (Sigma, St. Louis, MO) was added fresh during the media change. The media was then replaced with FAO assay medium containing 111mM NaCl, 4.7mM KCl, 1.25mM CaCl_2_, 2mM MgSO4,1.2mM NaH_2_PO_4_ supplemented with 2.5mM glucose, 0.5mM Carnitine and 5mM HEPES and incubated for 30-45 minutes. XF Palmitate BSA or BSA control was added to the medium just prior to the start of the XF assay. The OCR (pmol of O_2_/min) was measured after sequential addition of electron transport chain modulators Oligomycin, FCCP and Rotenone/Antimycin A.

### 2.11. Electron microscopy

Alveolar macrophages were purified by plating BAL fluid collected from WT and *Csf2rb*^*−/−*^ mice in 6 well plates for 2-4 hours in complete culture medium. The adherent cells were washed with prewarmed DPBS and macrophages were collected using TryPLE + 0.3% pluronic acid for 30-40 minutes at 37°C. Live cells were counted in a hemocytometer using trypan blue. Approximately, 1-2 million cells were harvested and centrifuged at 3000 rpm for 3 min. The cell pellet was resuspended in fresh Karnovsky’s fixative (contains 2.5% glutaraldehyde, 2% paraformaldehyde, 0.1M sodium cacodylate buffer, pH 7.3. Final 0.1% Cacl_2_) and submitted to the CCHMC Electron Microscopy core for further processing for transmission electron microscopy (TEM). The samples were post fixed in 1% osmium tetroxide prepared in 0.15M sodium cacodylate buffer, suspended in 1.5% agarose, processed through a graded series of alcohol solutions, followed by infiltration, and finally embedded in LX-112 resin. After polymerization at 60°C for 3 days, ultrathin sections (120nm) were cut using a Leica EM UC6 ultramicrotome and counterstained in 2% aqueous uranyl acetate and Reynold’s lead citrate. Images were taken using a transmission electron microscope (Hitachi H-7650, Tokyo, Japan).

### 2.12. Nuclear magnetic resonance (NMR) analysis

*Media collection* BMDMs from WT and *Csf2rb*^*−/−*^ mice were generated and cultured for NMR in complete culture medium containing 10% dialyzed serum (Sigma, St. Louis, USA) and with no HEPES addition. BMDMs (2×10^6^) were plated in a 100mm tissue culture dish with complete culture medium with dialyzed serum for 48 hours at 37C. This media is termed as conditioned media. Media was also collected at an initial time point (T_initial_ or T_0_ media) before culture. This media is termed “unused media”. Media (1 ml) was centrifuged at 2000g at 4°C for 5 minutes and the supernatant (800μl) was transferred into a new eppendorf tube, frozen in liquid nitrogen, and stored at −80°C. Just prior to the NMR, samples were thawed on ice and centrifuged 4000x *g*_*n*_ for 5 min at 4 °C. The supernatant (600 μL) were aliquoted onto pre-washed 3 kDa spin filters (NANOSEP 3K, Pall Life Sciences), centrifuged 10000x *g*_*n*_ for 90 min at 4 °C, and the media filtrate was mixed with NMR buffer up to 600 uL.

#### 2.12.1. Cell Sample Preparation

Adherent macrophages were washed twice with ice-cold PBS and cell extracts and protein were precipitated by adding cold MeOH/H2O (80:20). The macrophages were scraped vigorously, and the cell extracts were collected in 2ml Eppendorf tubes. The culture dishes were washed with 500 ul of MeOH/H_2_O solution and the cell extracts were pooled. The tubes were vortexed, incubated on ice for 5 minutes, centrifuged at 5,725xg for 5 minutes, and the supernatant was snap frozen for NMR analysis. The cell pellet was also snap frozen and stored for protein analysis via the manufacturer’s instructions for the Pierce BCA Protein assay kit (Thermo Fisher Scientific) to normalize the NMR data. Prior to the NMR analysis, the methanol phase was dried in a SpeedVac centrifuge for 4-6 h and stored at −20 °C. On the day of the NMR analysis, the dried polar extract samples were resuspended in 220 of NMR buffer (100 mM potassium phosphate (pH 7.3), 0.1% sodium azide, 1mM trimethylsilylproprionate (TSP) in 100% D_2_O).

#### 2.12.2 NMR Spectroscopy and data analysis

The experiments were conducted using 200 μL cell extracts samples in 103.5 mm × 3 mm NMR tubes (Bruker Analytik, Rheinstetten, Germany) and 550 μL media samples in 103.5 mm × 5 mm NMR tubes. The data was acquired on a Bruker Avance II 600 MHz spectrometer with Prodigy BBO cryoprobe as previously described (Maki et al., 2020). Metabolites were assigned based on 1D ^1^H and 2D NMR experiments by comparing the chemical shifts with reference spectra found in databases, such as the Human Metabolome Database (HMDB) (Wishart et al., 2007), the Madison metabolomics consortium database (MMCD) (Cui et al., 2008), the biological magnetic resonance data bank (BMRB) (Ulrich et al., 2008), and Chenomx^®^ NMR Suite profiling software (Chenomx Inc. version 8.1).

The quantification of metabolites was performed on Chenomx software based on the Trimethylsilylpropanoic acid (TMSP) internal reference in the NMR spectrum nuclear magnetic resonance of aqueous solvents. The cellular metabolites concentrations were normalized to total protein extracted from each sample to normalize for the differences in number of cells per sample. Normalized metabolites concentrations were then assessed by multivariate and univariate analysis. All the multivariate analysis were performed with MetaboAnalyst 4.0 (Chong et al., 2019) and univariate comparisons with RStudio.

### 2.13. Flow cytometry measurements

BMDMs were immunostained to detect Glut1, Glut3 (Novus biologicals) and CD36 (Thermofisher scientific) expression according to manufacturer’s instructions and evaluated using FACS Canto II.

### 2.14. Transcriptome Analysis of bone marrow-derived macrophages

Total RNA was isolated from alveolar macrophages using TRIzol Reagent (Life Technologies, Carlsbad, CA) according to manufacturer’s instructions. Total RNA (500 to 1000 ng) determined by Invitrogen Qbit high sensitivity spectrofluorometric measurement is poly A selected and reverse transcribed using Illumina’s TruSeq stranded mRNA library preparation kit. Each sample can be fitted with one of 24 adapters containing a different 8 base molecular barcode for high level multiplexing. After 15 cycles of PCR amplification, completed libraries are sequenced on an Illumina HiSeq2500 in Rapid Mode, generating 20 million or more high quality 75 base long paired end reads per sample.

### 2.15. Statistical Analysis

Numeric data were evaluated for a normal distribution using the Shapiro-Wilk and presented as mean ± SD for each group. Group comparisons were made using a two-tailed, non-parametric Mann-Whitney test. Statistical analysis was performed using GraphPad Prism 7 software. P-values of <0.05 were considered as statistically significant. All the experiments were repeated at least 3 times.

## 3.0. Results

### 3.1. GM-CSF stimulates cell cycle progression and proliferation of macrophages

Because bone marrow-derived macrophages (BMDMs) proliferate in proportion to endogenous pulmonary GM-CSF concentration after pulmonary transplantation into *Csf2rb* deficient (*Csf2rb*^*−/−*^) mice (Arumugam et al., 2019; Suzuki et al., 2014), we first determined the cell-cycle status (i.e., G0/G1→S→G2→M) of bone marrow-derived macrophages *in vitro.* Compared to wild type (WT) BMDMs, *Csf2rb*^*−/−*^ BMDMs had significantly reduced cell-cycle progression (Fig. 1A, B). Next, we measured cellular proliferation of BMDMs using an assay based on quantification of mitochondrial metabolic activity-based formazan chromophore formation, which showed that *Csf2rb*^*−/−*^ BMDMs had reduced proliferation compared to WT BMDMs (Fig. 1C). Since cell-cycle progression and proliferation require energy, we evaluated the metabolic activity of BMDMs, which showed that *Csf2rb*^*−/−*^ BMDMs had markedly reduced metabolism compared to WT at all time points evaluated (Fig. 1D). Since the production of reactive oxygen species (ROS) is proportional to cellular metabolism and proliferation, we measured the effects of GM-CSF deprivation on production of singlet oxygen (^1^O_2_), superoxide (^·^O_2_), and hydroxyl radical (HO^·^) generation using a DCFDA-based assay, which showed ROS production by *Csf2rb*^*−/−*^ BMDMs was significantly reduced compared to WT BMDMs (Fig. 1E). Together, these results show that the metabolic activity of macrophages was reduced in the absence of GM-CSF stimulation and correlated with reduced cellular proliferation.

**Fig. 1.**
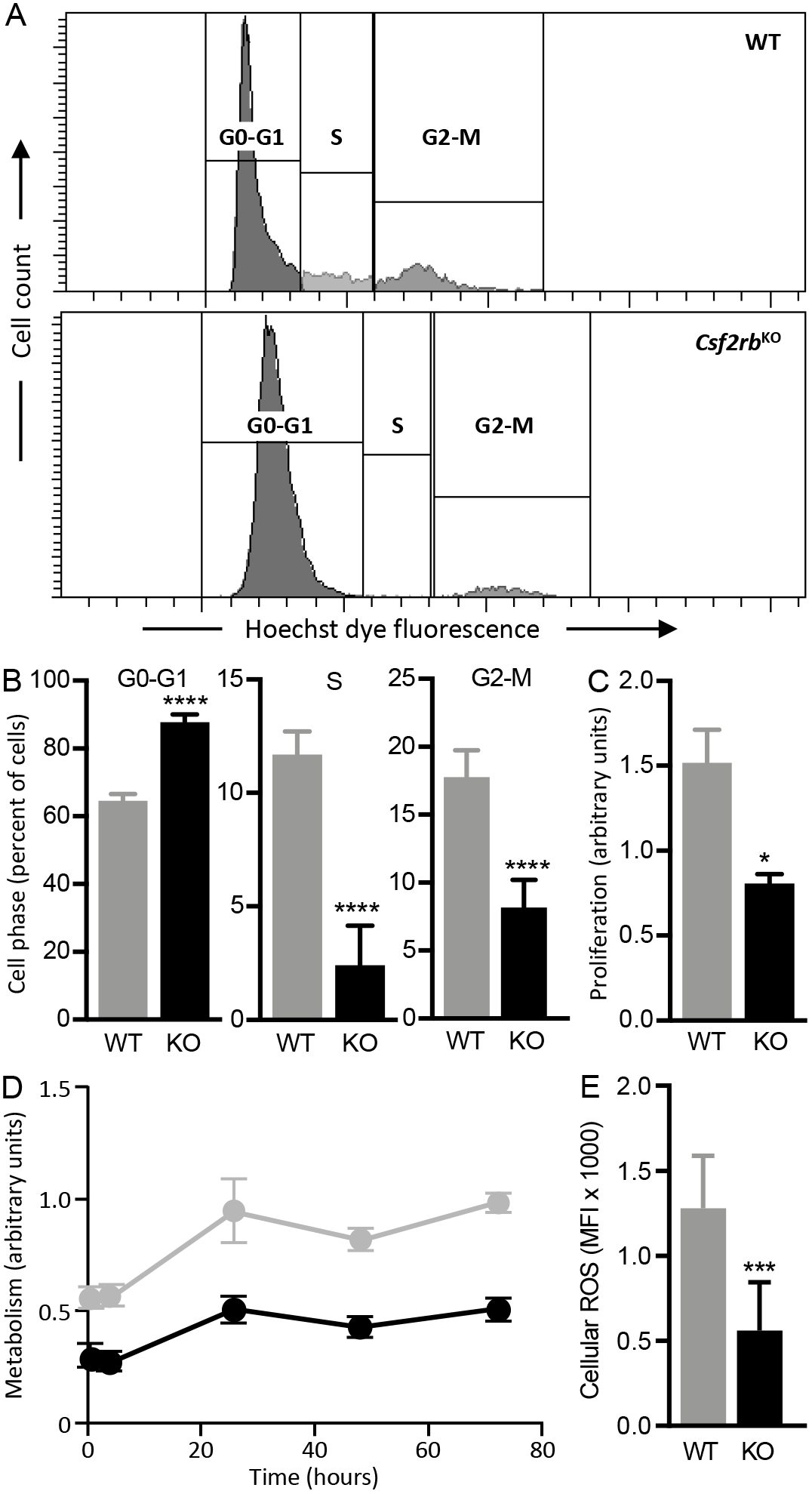
GM-CSF induces cell cycle progression and macrophage proliferation. (A) Cell cycle analysis in WT and *Csf2rb−/−* BMDMs (n=6-8 samples/group) using Hoechst staining. Representative flow cytometer plots showing GO (gap 0)-Gl (gap 1), S phase (DNA synthesis) and G2 (gap 2)-M (mitosis) phases of the cell cycle. (B) Quantification of the percentage of cells present in different phases of cell cycle. (C) WST-1 cell proliferation assay in BMDMs, to measure of mitochondrial dehydrogenase activity in metabolically active cells. (D) MTT cell proliferation assay that measures the reducing potential of cells. (E) Mean fluorescence intensity (MFI) of DCF fluorescence to measure intracellular ROS. *P<0.05, ***P<001, ****P<0.0001.

### 3.2. GM-CSF is required for Mitochondrial Mass, Integrity, and Function in Macrophages

Since our data revealed decrease in the metabolic activity in the absence of GM-CSF signaling, we evaluated the effects of GM-CSF on mitochondrial structure and function. AMs from WT or *Csf2rb*^*−/−*^ mice were immunostained with an antibody to Translocase of Outer Mitochondrial Membrane 20 (TOMM20), an integral membrane protein distributed across the outer mitochondrial membrane (OMM) surface, as well as with wheat germ agglutinin (to label the macrophage plasma membrane), and 4’,6-diamidino-2-phenylindole (to label the macrophage nucleus) and examined by confocal immunofluorescence microscopy. TOMM20 was chosen instead of MitoTracker green chromophore because binding of the former is unaffected by mitochondrial activity and membrane potential (de Almeida et al., 2017; Ito and Suda, 2014; Keij et al., 2000). Results show that mitochondrial mass was higher in *Csf2rb*^*−/−*^ macrophages compared to WT macrophages (Fig. 2A). Quantification of TOMM20 fluorescence showed that, compared to WT macrophages, *Csf2rb*^*−/−*^ macrophages had a 2.5-fold higher mean fluorescence intensity indicating an increase in mitochondrial mass (Fig. 2B).

**Fig. 2.**
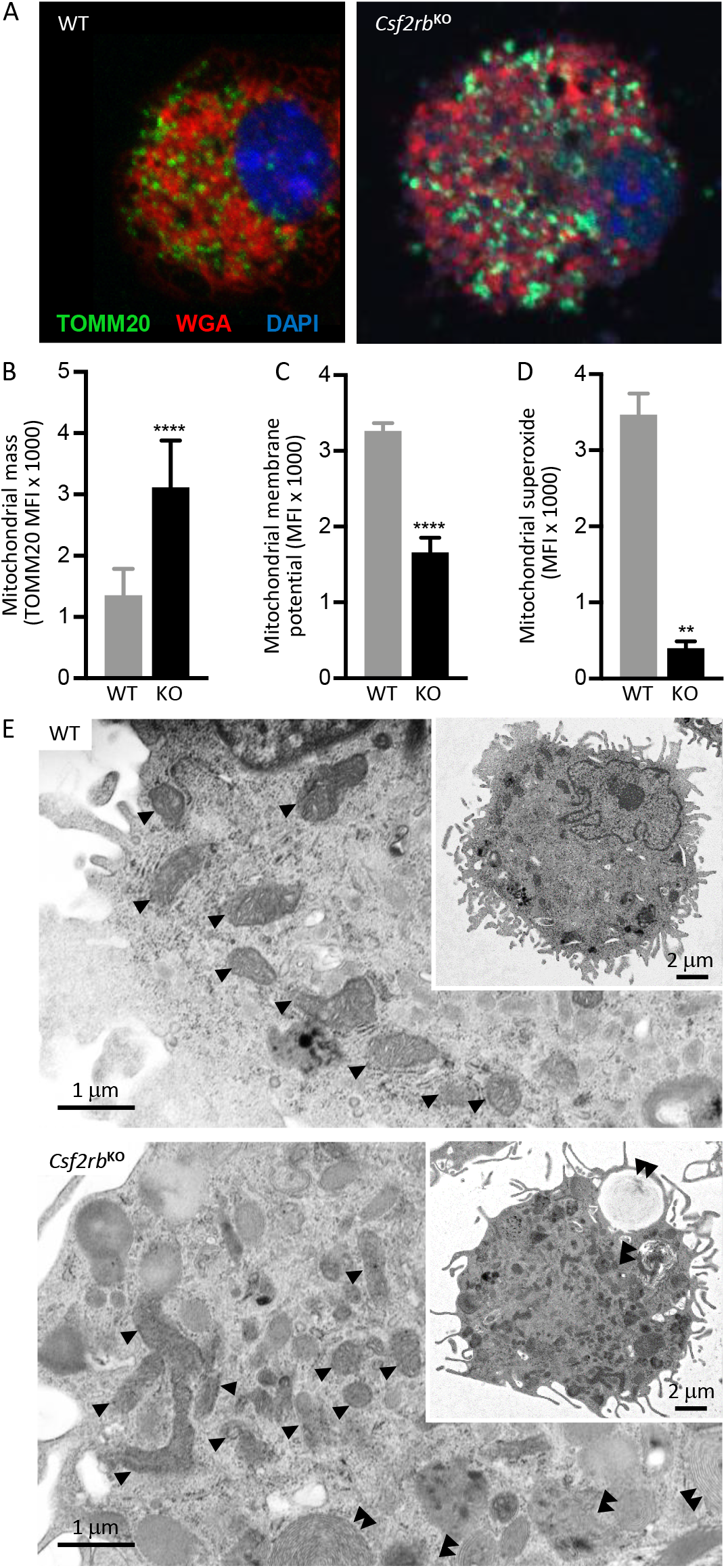
Absence of GM-CSF signaling results in accumulation of dysfunctional mitochondria. (A) Immunofluorescence images of alveolar macrophages from WT and *Csf2rb*^*−/−*^ macrophages stained with DAPI (blue) WGM-(red), TOMM20 (green) are shown. (B) Quantification of mean fluorescent intensity of TOMM20 staining by Nikon Imaging software (NIS) elements software to measure mitochondrial mass. (C) Quantification of MFI of TMRE fluorescence to assess mitochondrial membrane potential by flow cytometry. (D) Quantification of MFI of red fluorescence produced in WT and *Csf2rb*^*−/−*^ macrophages due to oxidation of the reagent MitoSox red by mitochondrial superoxide generated in the cells. (E) Representative transmission electron micrograph images showing increased number of abnormally shaped mitochondria in *Csf2rb*^*−/−*^ macrophages compared to WT macrophages. Single black arrowheads point to mitochondria in WT and *Csf2rb*^*−/−*^ macrophages. Double arrow heads represent presence of inclusion and lamellar bodies in *Csf2rb*^*−/−*^ macrophages. Scale bar represents 1μM and 2 μM (inset) software. **P<0.01, ****P<0.0001.

Next, mitochondrial membrane potential (MMP) was measured using tetramethylrhodamine ethyl ester (TMRE), a cell-permeant, cationic fluorescent dye readily sequestered by active mitochondria due to their negative charge on the outer membrane surface. Results showed that TMRE uptake was reduced in *Csf2rb*^*−/−*^ macrophages compared to WT macrophages (Fig 2C), consistent with depolarization or inactivation of mitochondria.

Production of mitochondrial superoxide species 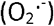 was also measured using MitoSox Red reagent, a fluorogenic molecule specifically targeted to mitochondria in live cells that fluoresces in proportion to superoxide 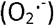 production. Mitochondrial superoxide production was reduced in *Csf2rb*^*−/−*^ macrophages compared to WT macrophages (Fig. 2D).

Since *Csf2rb*^*−/−*^ macrophages had an increased mitochondrial mass with reduced membrane potential and reduced ROS production, we evaluated mitochondrial ultrastructure using transmission electron microscopy. Results confirmed the increase in mitochondrial mass and identified ultrastructural abnormalities including increased mitochondrial numbers of large, hyperfused, and also short mitochondria in *Csf2rb*^*−/−*^ macrophages compared to WT macrophages (Fig. 2E).

Together, these results indicate that GM-CSF signaling is required for maintenance of mitochondrial structure, integrity and functions in macrophages.

### 3.3. GM-CSF Regulation of Fatty Acid-Based Energy Production in Macrophages

To maintain lung homeostasis, AMs must constitutively clear fatty acid-containing surfactant phospholipids – a function that requires GM-CSF stimulation. Since fatty acid catabolism occurs by β-oxidation in mitochondria and expression of fatty acid transporters CPT1a and CPT2 (located in the outer and inner mitochondrial membrane, respectively) also require GM-CSF stimulation, we examined the effects of GM-CSF on mitochondrial metabolism in macrophage by measuring OCR using a Seahorse extracellular flux analyzer with the XF Long Chain Fatty Acid Oxidation Stress Test Kit using palmitate as the exogenous fuel source. Cells were cultured in substrate-limited medium supplemented with carnitine to prime them to metabolize exogenous fatty acids. Compared to WT macrophages, *Csf2rb*^*−/−*^ macrophages had significant reductions in basal respiration, ATP-linked respiration, maximal respiration, and non-mitochondrial respiration (Fig. 3A, B). Under these conditions, the spare respiratory capacity of *Csf2rb*^*−/−*^ macrophages, a measure of the cellular capacity to respond to an energy demand, was zero, indicating an absence of the ability to respond to an energy demand through β-oxidation of palmitate. To evaluate the potential for a cellular response to reduced fatty acid oxidation in the absence of GM-CSF signaling, we measured expression of CD36, which participates in cellular uptake of long-chain fatty acids. Results show it was significantly upregulated in *Csf2rb*^*−/−*^ compared to WT macrophages (Fig. 3C), indicating that macrophages were unable to metabolize exogenous free fatty acids for energy production in the absence of GM-CSF stimulation.

**Fig. 3.**
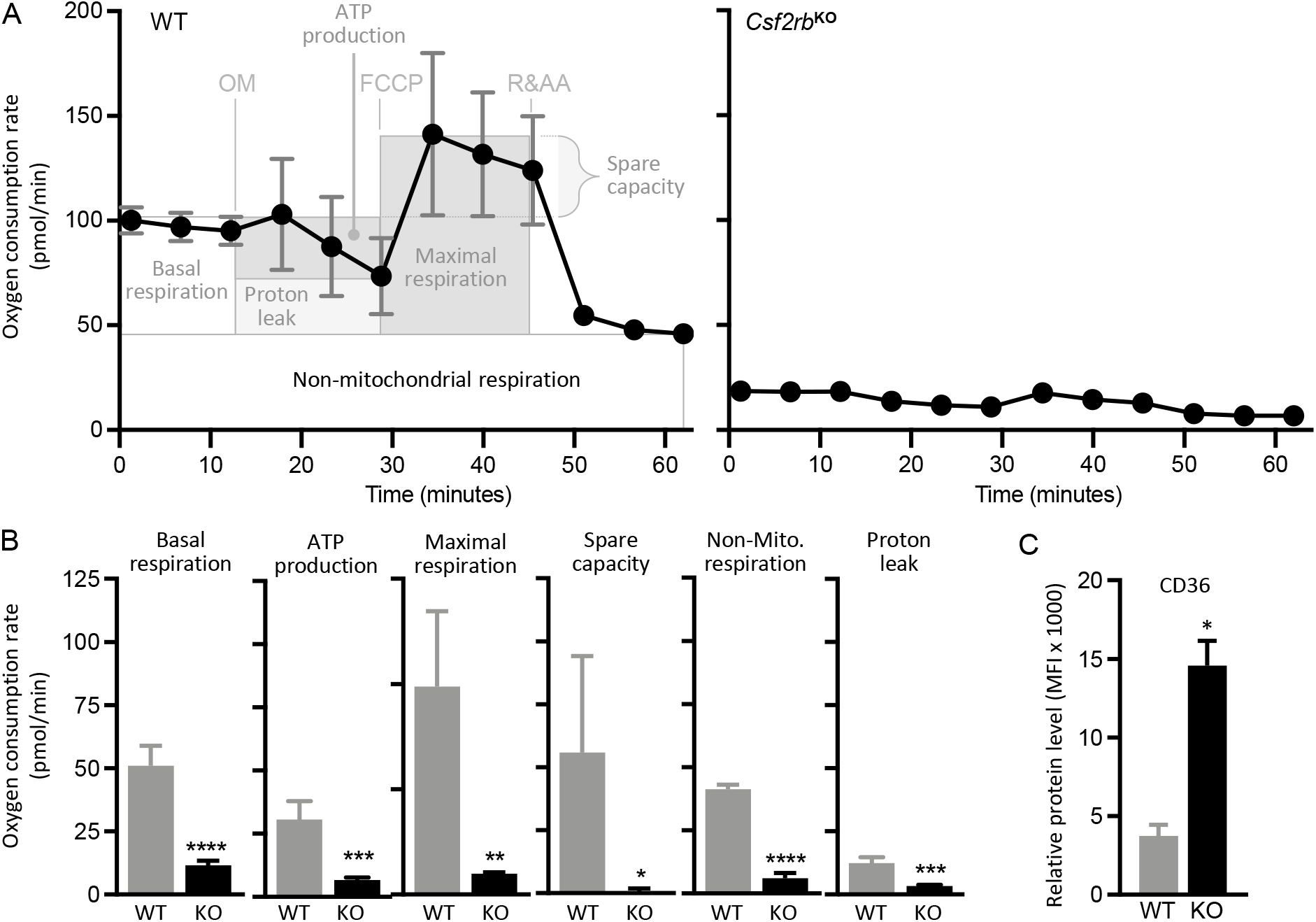
Fatty acid oxidation in macrophages impaired in the absence of GM-CSF signaling. BMDMs from WT and *Csf2rb*^*−/−*^ mice (n=6-8 mice/group) were incubated in substrate limited medium in the presence of palmitoyl and carnitine and OCR measurements were done. (A) Representative profile of XF Palmitate oxidation stress test was used to measure changes in OCR in BMDM of WT and *Csf2rb*^*−/−*^ mice at baseline, in the presence of ATP synthase inhibitor oligomycin (OM), after addition of uncoupler of oxidative phosphorylation (FCCP, carbonyl cyanide-4-(trifluoromethoxy) phenylhydrazone) and inhibitors of mitochondrial respiratory chain complexes I (Rotenone, (R)) and III (Antimycin A, (AA)). (B) Quantification of respiration parameters including basal respiration, maximal respiration, ATP production, spare respiratory capacity, non-mitochondrial respiration, and proton leak. (C) CD36 expression on macrophage cell surface measured by flow cytometry. Relative CD36 expression in WT and *Csf2rb*^*−/−*^ macrophages is represented as the mean fluorescence intensity. **P<0.01, ***P<0.001, ****P<0.0001.

### 3.4. GM-CSF Regulation of Glucose-Derived Energy Production in Macrophages

To better define the mechanism(s) by which GM-CSF regulates metabolism in macrophages, we measured cellular oxygen consumption rate (OCR) – a correlate of oxidative phosphorylation, and extracellular acidification rate (ECAR) – a correlate of glycolysis, using a Seahorse extracellular flux analyzer with a XF Cell Mito Stress Test Kit and Glycolysis Stress Test Kit, respectively, using glucose as the exogenous fuel source. Compared to WT macrophages, *Csf2rb*^*−/−*^ macrophages had significant reductions in basal respiration, ATP-linked respiration, maximal respiration, and non-mitochondrial respiration (Fig, 4A, C). Furthermore, glycolysis and glycolytic capacity (maximum conversion of glucose to pyruvate, which correlates with ECAR) were significantly reduced in *Csf2rb*^*−/−*^ macrophages compared to WT macrophages (Fig. 4B, D). In addition, the glycolytic reserve (a measure of the cellular capacity to respond to an energy demand as well as the proximity of its glycolytic functional capacity to the theoretical maximum) was very small in both WT and *Csf2rb*^*−/−*^ macrophages (Fig. 4B, D). Overall, these results demonstrate that GM-CSF regulates glucose metabolism in macrophages to generate energy needed for cell survival and proliferation.

**Fig. 4.**
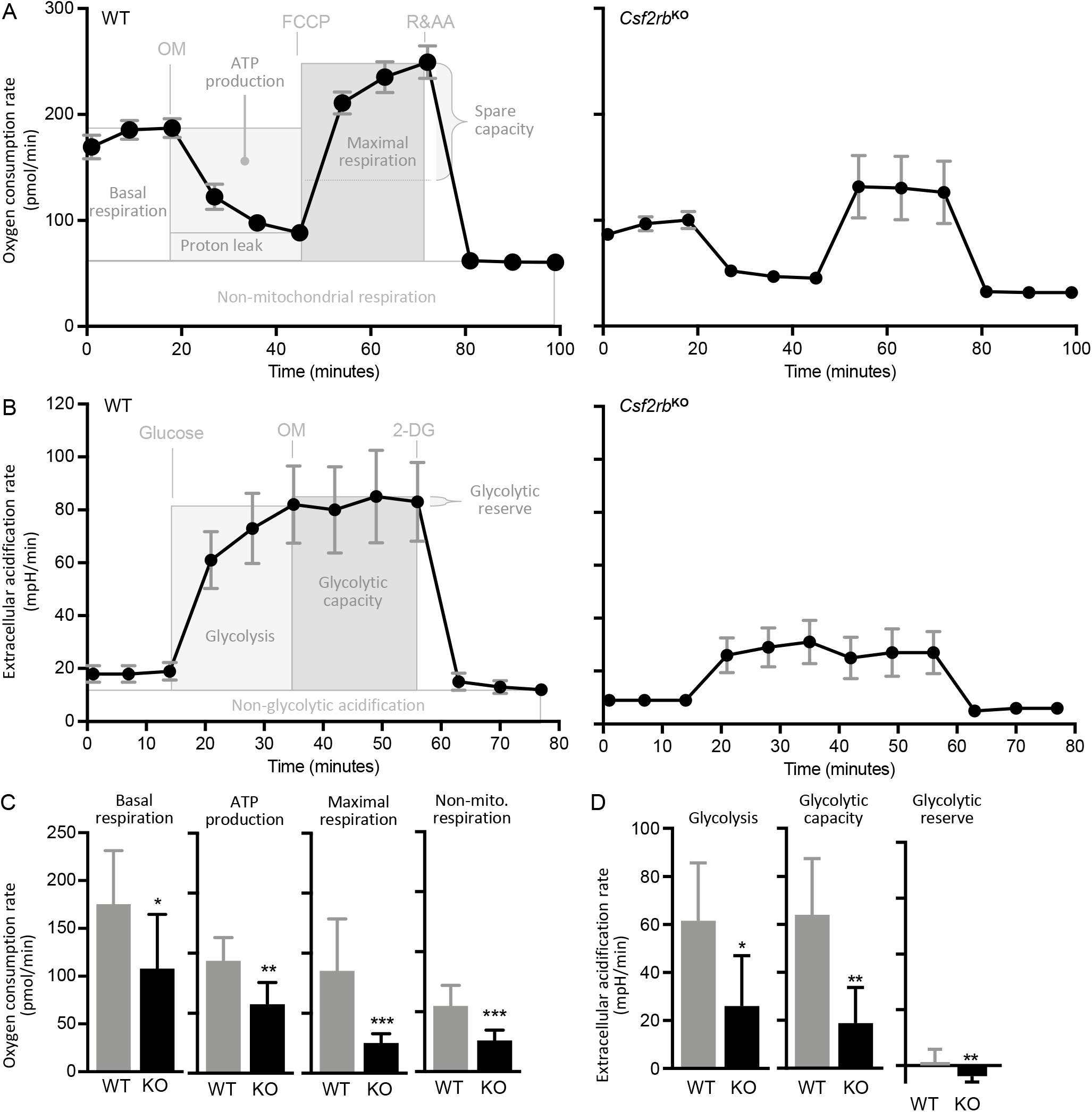
GM-CSF signaling stimulates glycolysis and mitochondrial oxidative phosphorylation in macrophages. (A) Representative profiles of mitochondrial stress test to measure oxygen consumption rate (OCR) in BMDMs of WT and *Csf2rb*^*−/−*^ mice (n=8 mice/group) at baseline, in the presence of the ATP synthase inhibitor, oligomycin (OM), after addition of an uncoupler of oxidative phosphorylation (FCCP, carbonyl cyanide-4-(trifluoromethoxy phenylhydrazone), and in the presences of inhibitors of mitochondrial respiratory chain complexes I (Rotenone; (R)) and III (Antimycin A (AA)). (B) Representation profiles of extracellular acidification rate (ECAR) measurements as an indicator of glycolysis. ECAR measurements were done by sequential addition of glucose, oligomycin (OM) and 2-deoxyglucose (2-DG, an inhibitor of hexokinase-2 catalyzing the first step in glycolysis). (C) Quantification of respiration parameters including basal respiration, maximal respiration ATP production and non-mitochondrial respiration. (D) Quantification of glycolysis, glycolytic capacity, and glycolytic reserve. All error bars represent mean ± SEM.

### 3.5. GM-CSF Regulates Multiple Metabolic Pathways Required for Macrophage Proliferation

To broadly address the potential role of GM-CSF in macrophage metabolism, NMR-based metabolomics were performed on cultured BMDMs from WT and *Csf2rb*^*−/−*^ mice and on conditioned media (CM) to simultaneously quantify biochemical substrates, intermediates, and byproducts of several metabolic pathways including glycolysis, the tricarboxylic acid (TCA) cycle, the pentose phosphate pathway, and amino acid biosynthesis. One-dimensional ^1^H spectral analysis of extracts from cells or CM (following *in vitro* culture for 48 hours) identified 39 or 28 differentially expressed metabolites from WT or *Csf2rb*^*−/−*^ BMDMs, respectively (Figure 5). A principal component analysis scatterplot of the results revealed strikingly different profiles for intracellular metabolites in WT and *Csf2rb*^*−/−*^ BMDMs with principal component 1 (PC1) and PC2 accounting for 82.3% and 10.9% of the variance, respectively (Fig. 5A).

**Fig. 5.**
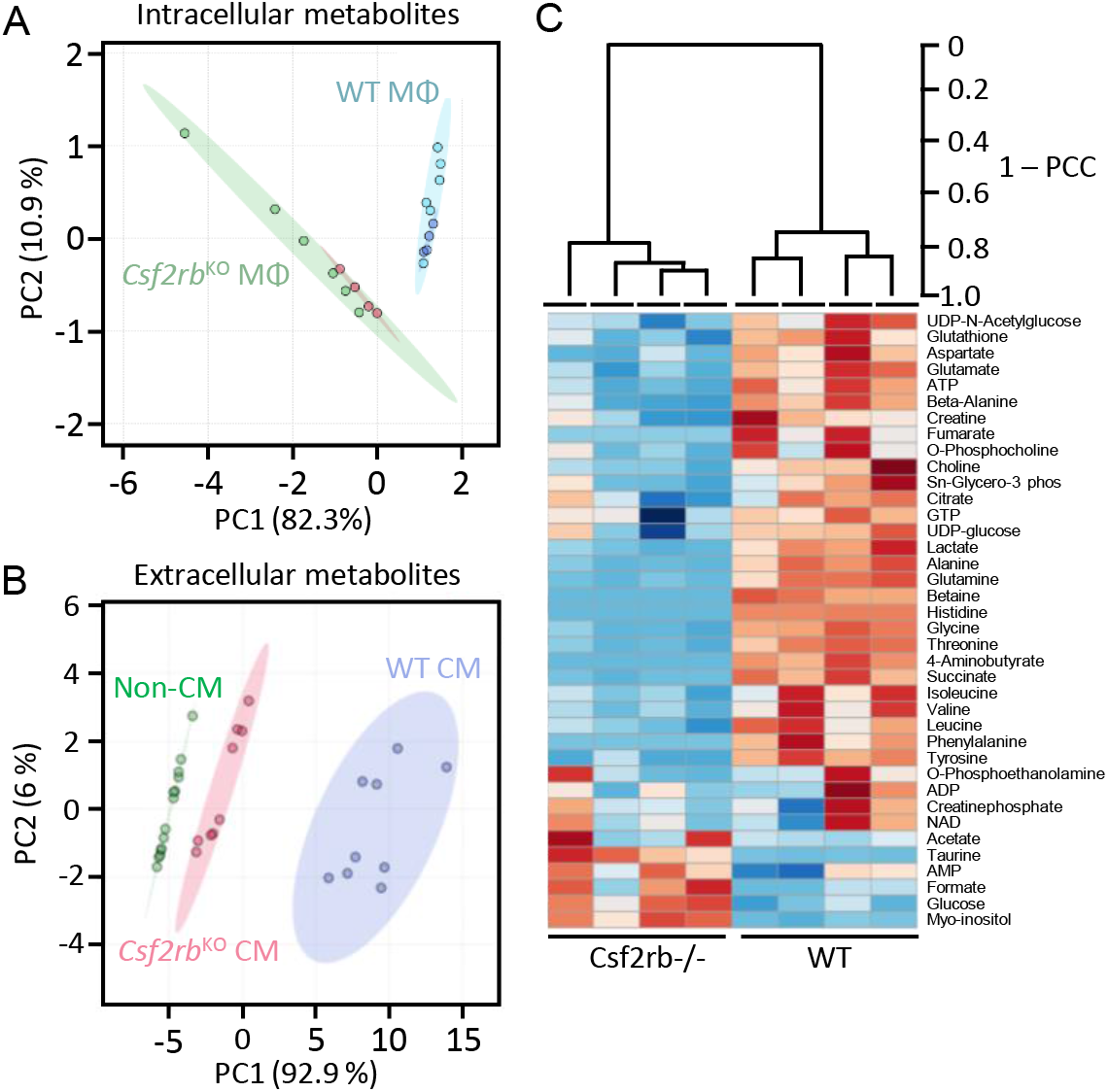
Macrophage metabolite profile measured by NMR metabolomics approach reveal differences in the intracellular and extracellular metabolite profiles between WT and *Csf2rb*^*−/−*^ macrophages (n=9-10/group). Two-dimensional principal component analysis generated from (A) intracellular and (B) extracellular (non-conditioned and conditioned media) metabolite profiles is shown. (C) Hierarchical clustering analysis and representative heat map visualization of 38 metabolites detected from WT and *Csf2rb*^*−/−*^ macrophages (n=4/group). Rows represent metabolites and columns indicate WT and *CsfZrb*^*−/−*^ samples. Red indicate metabolites detected in high abundance and blue represents metabolites detected in low abundance.

Similarly, the results for extracellular metabolites also revealed markedly different profiles in WT and *Csf2rb*^*−/−*^-CM with PC1 and PC2 accounting for 92.9% and 6% of the total variance, respectively (Fig.5B). Further, the scatterplot profile of metabolites in *Csf2rb*^*−/−*^-CM closely resembled that of unconditioned media but was widely separated from that of WT-CM (Fig. 5B). Unsupervised hierarchical clustering and heat map analysis of BMDM extracts demonstrated widely differing segregation of the metabolic profiles between WT and *Csf2rb*^*−/−*^ BMDMs and similar profiles of the respective strain-specific replicates (Fig. 5C).

#### 3.5.1. Glycolysis

We first examined biomolecules involved in glycolysis, the primary energy producing metabolic pathway in the cytoplasm. Compared to WT macrophages, the intracellular glucose concentration was significantly increased while lactate, UDP-glucose, and UDP-N-acetyl glucosamine concentrations were significantly decreased in *Csf2rb*^*−/−*^ BMDMs, suggesting glycolytic metabolism was reduced in the absence of GM-CSF signaling (Fig. 6A). Consistent with this possibility, the extracellular glucose concentration was reduced to a lesser degree in *Csf2rb*^*−/−*^ CM than in WT-CM (both compared to unconditioned media) while the cell-derived lactate concentration was increased to a lesser degree in *Csf2rb*^*−/−*^-CM than in WT-CM (both compared to unconditioned media) (Fig. 6B). Further, expression of the major glucose transporters (Glut1 and Glut3) normally present in macrophages were increased in *Csf2rb*^*−/−*^ BMDMs compared to WT BMDMs (Figure 3C). Together, these findings support the conclusion that glycolysis is reduced in macrophages in the absence of stimulation by GM-CSF.

**Fig. 6.**
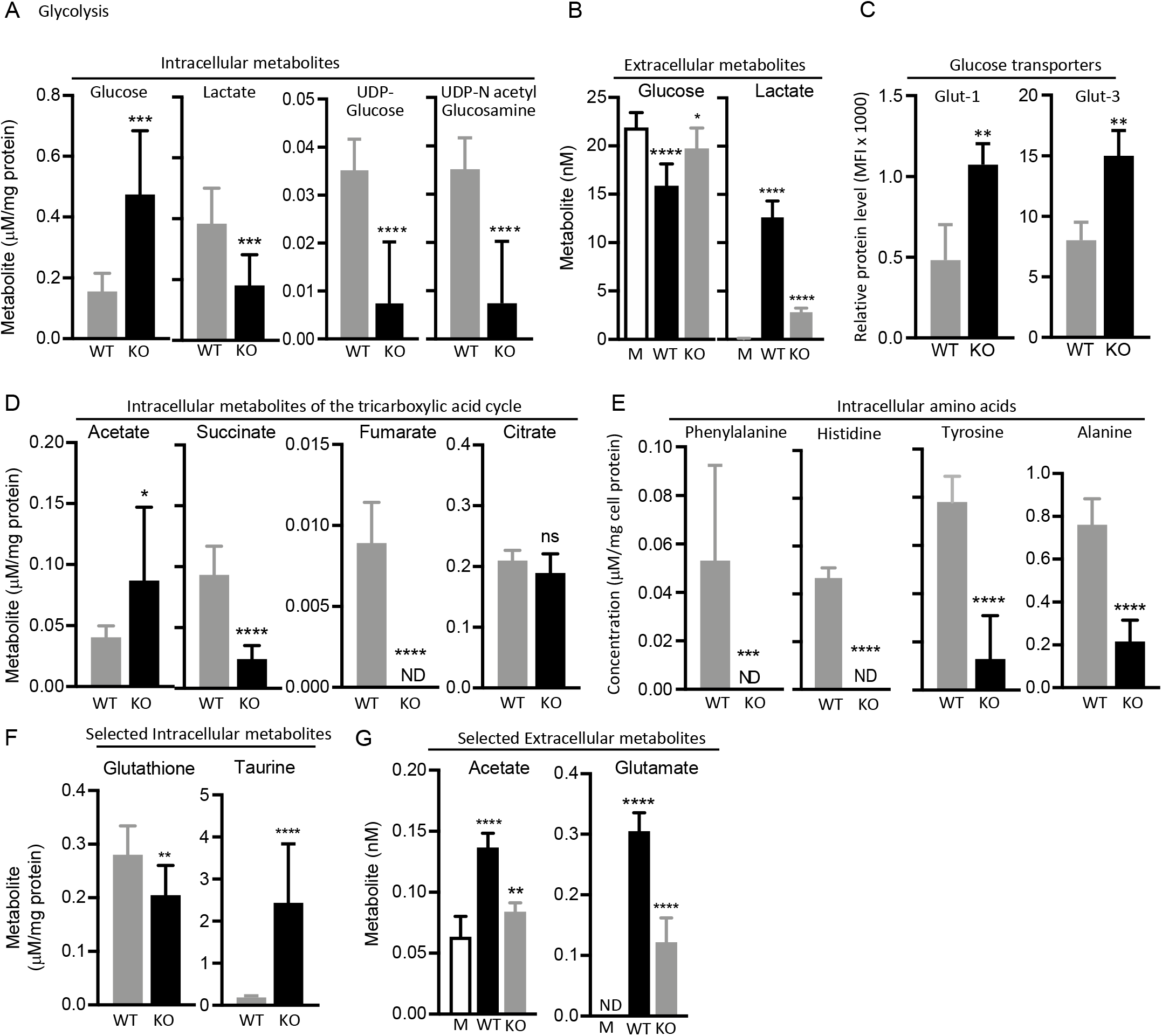
GM-CSF regulates metabolic pathways for macrophage proliferation, levels of (A) intracellular metabolites of glycolysis (glucose, lactate) and glycolytic intermediate derivatives UDP glucose and UDP-N acetyl glucosamine, (B) extracellular metabolites, including glucose and lactate, (D) tri-carboxylic acid metabolites, acetate, succinate, fumarate, and citrate, (E) intracellular amino acids, phenylalanine, tyrosine, histidine and alanine, (F) intracellular metabolites glutathione and taurine and (G) extracellular metabolites acetate and glutamate were quantitated via NMR metabolomics measurements, (C) Level of glucose transporter (Glut-1 and Glut-3) expression measured by flow cytometry. Abbreviations: Glut-1, Glucose transporter 1; Glut-3, Glucose transporter, ns - not significant, *P<0.05, **P<0.01, ***P<0.001, ****P<0.0001.

#### 3.5.2. TCA Cycle

Next, we examined metabolites related to the TCA cycle, a series of chemical reactions in mitochondria that produce energy through oxidation of acetyl coenzyme A (acetyl-CoA) derived from carbohydrates, fats, and proteins. Compared to WT macrophages, the intracellular concentrations of succinate and fumarate were significantly decreased (the latter was undetectable) and acetate was significantly increased in *Csf2rb*^*−/−*^ macrophages compared to WT macrophages; citrate levels were similar (Fig. 6D). Together, the reduction in key TCA cycle intermediates coupled with increased acetate (which is consumed in converting pyruvate to acetyl-CoA a principal metabolite consumed in TCA cycle) suggest decreased mitochondrial regulation. The increase in acetate levels could be due to *de novo* acetate production, which can occur in limited metabolic environments such as during mitochondrial dysfunction to maintain acetyl-CoA pools and cell proliferation (Liu et al., 2018). These data suggest that TCA cycle-based metabolism is reduced in macrophages in the absence of GM-CSF stimulation.

#### 3.5.3. Pentose Phosphate Pathway and Amino Acid biosynthesis

The pentose phosphate pathway (also called the hexose monophosphate shunt or phospho-gluconate pathway) is a cytosolic pathway parallel to glycolysis that generates NADPH, pentoses, and ribose 5-phosphate, which are used in the synthesis of fatty acids, amino acids, and nucleotides, respectively. Compared to WT macrophages, the intracellular concentrations of amino acids normally synthesized from glycolysis, the TCA cycle, and pentose phosphate pathway were markedly altered in *Csf2rb*^*−/−*^ macrophages. Importantly, the levels of aromatic amino acids, phenylalanine and histidine were undetectable (Fig. 6E), while the levels of other amino acids (alanine, tyrosine, glycine, leucine, valine, glutamine, aspartate, threonine, and isoleucine) were significantly reduced (Fig. 6E, Fig. S1A). Thus, these protein building blocks, and sources of energy used for survival in starved cells were reduced in *Csf2rb*^*−/−*^ macrophages. The reduced extracellular concentration of glutamate (Fig. 6G) was consistent with these results, which together, suggest decreased synthesis/secretion of non-essential amino acids in the absence of GM-CSF.

#### 3.5.4. Oxidation/Reduction Metabolites

Glutathione, an important regulator of cell proliferation, was significantly reduced in *Csf2rb^−/−^* macrophages compared to WT macrophages (Fig. 6F). Glutathione is synthesized in the cytosol from L-glutamate and L-glycine, both of which were significantly reduced in *Csf2rb−/−* macrophages compared to WT macrophages (Fig. 6G, Fig. S1). Taurine, an important regulator of the cellular redox homeostasis, (Seidel et al., 2019) was significantly increased in *Csf2rb*^*−/−*^ compared to WT macrophages (Fig. 6F), suggesting accumulation and reduced metabolism leading to the observed decrease in ROS production (Fig. 1E) through its effects on mitochondrial bioenergetics. Since the glutathione-redox system utilizes electrons from NADPH to maintain ribonucleotide reductase activity, a rate-limiting enzyme in DNA synthesis, these results suggest that GM-CSF signaling deficiency may impact cellular proliferation by impairing DNA synthesis.

Together, these metabolomics results indicate that GM-CSF simulation has profound effects on several metabolic pathways relevant to cellular proliferation including cytosolic and mitochondrial metabolism.

### 3.6. Transcriptomic Analysis of GM-CSF-Regulated Involved in Macrophage Metabolism

To broadly evaluate GM-CSF regulation of genes important in cellular metabolism, genome-wide mRNA expression profiling was performed on WT and *Csf2rb*^*−/−*^ macrophages followed by supervised analyses of genes related to glycolysis, the TCA cycle, pentose phosphate pathway, and fatty acid oxidation.

#### 3.6.1. Glycolysis

Transcript levels of glycolysis-relevant enzymes were consistently altered among macrophage replicates from *Csf2rb*^*−/−*^ mice compared to WT mice (Fig. 7A). Importantly, transcripts encoding hexokinase isoform 2 (HK2), which catalyzes the essential initial step of glucose metabolism (conversion of glucose to glucose-6-phosphate) was consistently down-regulated in all replicates of *Csf2rb*^*−/−*^ macrophages compared to WT macrophages (Fig. 7A). Further, transcripts encoding ATP-dependent 6-phospho-fructokinase 2 *(PfKI)*, the enzyme that converts D-fructose 6-phosphate to fructose 1,6-bisphosphate, a key regulator, which commits glucose to the first step of glycolysis, and lactate dehydrogenase a (LDHa), which catalyzes the conversion of pyruvate to lactate, were both consistently reduced in *Csf2rb*^*−/−*^ macrophages compared to WT macrophages (Fig. 7A). Transcripts for other genes important to glycolysis *(Myc, Hif1a*, and *Ppargc1a)* were also decreased in *Csf2rb*^*−/−*^ macrophages compared to WT macrophages (Fig. 7A). Inhibition of this step prevents glucose turnover that produce NADH via glycolysis (Fig. 7A).

**Fig. 7.**
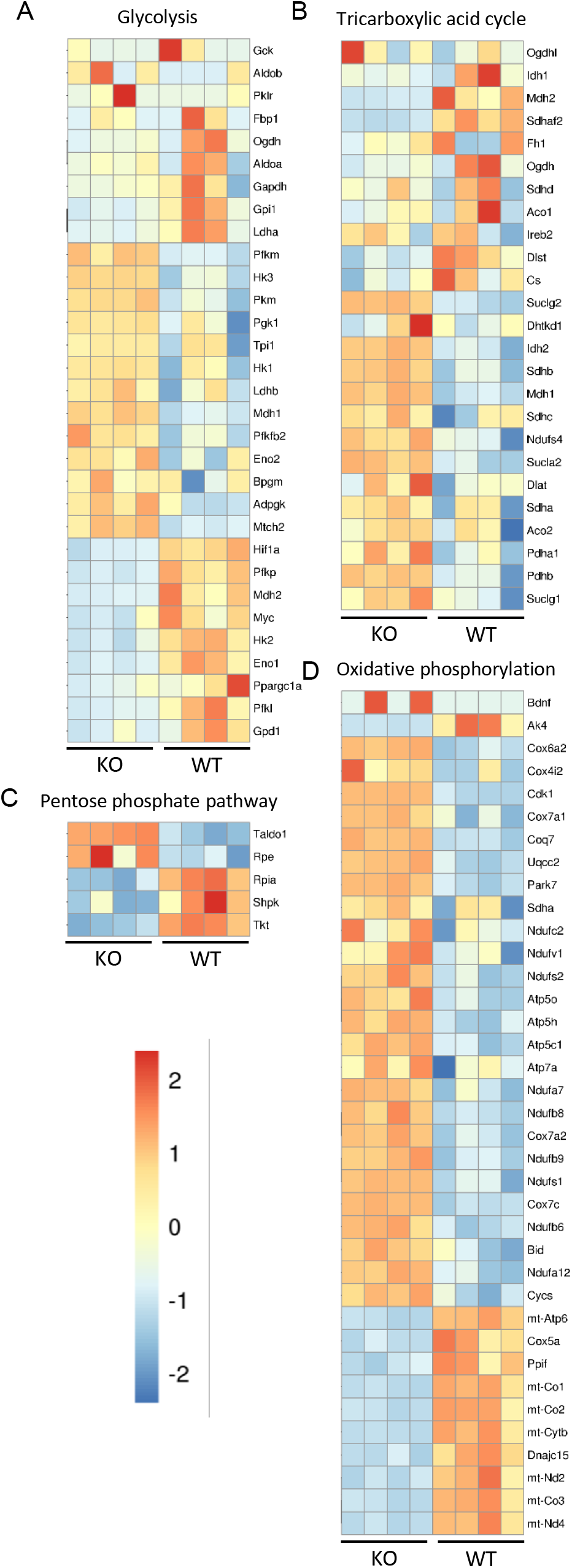
Gene expression profiles of metabolic pathway enzymes from WT and *CsfZrb*^*−/−*^ macrophages (n=4/group). Heat maps showing differential expression of genes involved in (A) glycolysis, (B) tricarboxylic acid cycle, (C) pentose phosphate pathway, and (D) oxidative phosphorylation are shown. Columns represent samples and rows represent genes. Color scale represents the abundance of gene transcripts. Genes with increased transcript levels are shown in red and decreased transcript levels in blue.

#### 3.6.2. TCA Cycle

Transcript levels for TCA cycle-relevant enzymes were also consistently altered among macrophage replicates from *Csf2rb*^*−/−*^ compared to WT mice (Fig. 7B). Specifically, transcripts for citrate synthase (Cs), the ubiquitous, rate-limiting enzyme that commits acetyl-CoA to the TCA cycle by combining with oxaloacetate and water to form citrate and reduced CoA, were consistently reduced in *Csf2rb*^*−/−*^ macrophages compared to WT macrophages (Fig. 7B). Further, transcripts for other key TCA cycle enzymes (mitochondrial malate dehydrogenase 2 (*Mdh2*), fumarate hydratase 1 (*Fh1*), succinate dehydrogenase assembly factor 2 *(Sdhaf2)*, succinate dehydrogenase complex and subunit D integral membrane protein (*Sdhd*)) were also reduced in *Csf2rb*^*−/−*^ macrophages compared to WT macrophages (Fig. 7B).

#### 3.6.3. Pentose phosphate pathway

The pentose phosphate pathway comprises an initial, oxidative phase in which NADPH is generated and a subsequent, non-oxidative phase in which 5-carbon sugars and ribose 5-phosphate are produced and used in the synthesis of amino acids and nucleotides, respectively. Transcript levels for pathway components in the oxidative phase were consistently increased while those in the non-oxidative phase were consistently decreased in *Csf2rb*^*−/−*^ macrophages compared to WT macrophages (Fig. 7C). Of particular relevance to DNA synthesis, transcript levels for non-oxidative or ‘anabolic’ phase genes (ribose 5-P04 isomerase A (*Rpia*), sedoheptulose kinase *(ShpK)* and transketolase *(tkt))* were consistently reduced in replicates of *Csf2rb*^*−/−*^ macrophages compared to WT macrophages (Fig. 7C), implying that conversion of glucose to ribose-5-phosphates for the synthesis of nucleotide precursors is reduced.

#### 3.6.4. Oxidative Phosphorylation

Energy production via oxidative phosphorylation involves three major steps: 1) oxidation-reduction reactions involving electron transfers between specialized proteins embedded in the inner mitochondrial membrane; 2) generation of a proton (and electrical) gradient across the inner mitochondrial membrane; and 3) synthesis of ATP using energy from the spontaneous diffusion of electrons down the proton gradient generated in step 2. Transcript levels encoding genes involved in oxidative phosphorylation were very consistently altered in *Csf2rb*^*−/−*^ macrophages compared to WT macrophages (Fig. 7D). Interestingly, transcripts for a number of mitochondrial encoded genes including cytochrome c oxidase – the terminal enzyme of the electron transport chain – (e.g., subunits I, II, and III *(Mt-co1, Mt-co2, Mt-co3)*, NADH dehydrogenase 2 and 4 *(Mt-Nd2, Mt-Nd4)*, cytochrome c oxidase subunit 4A *(Cox5a)*, cytochrome b *(Mt-Cytb)*, peptidylprolyl isomerase f *(Ppif)* and ATP synthase 6 *(Mt-Atp6)* were consistently and markedly reduced in *Csf2rb*^*−/−*^ macrophages compared to WT macrophages (Fig. 7D). In contrast, transcripts for non-mitochondrial genes encoding NADH dehydrogenase (ubiquinone) subunits (e.g., *Ndufa7, Ndufb8, Ndufb9, Nfduc2, Ndufs1, Ndufs2)* were consistently upregulated in *Csf2rb*^*−/−*^ macrophages compared to WT macrophages (Fig. 7D). Together, these transcriptomics results indicate that GM-CSF signaling is important for the expression of mitochondrial encoded genes relevant to ATP production and cytochrome c-dependent apoptosis.

### 3.7. GM-CSF Deficiency Impairs ATP Production in Macrophages

Since several lines of evidence indicate that disruption of GM-CSF signaling in macrophages results in reduced ATP generation, we measured the intracellular levels of ATP and its precursors, ADP and AMP, using NMR. Compared to WT macrophages, *Csf2rb*^*−/−*^ macrophages had reduced levels of ATP and increased intracellular levels of AMP and ADP (Fig. 8A), resulting in increased ratios of AMP: ATP, ADP:ATP and AMP:ADP (Fig. 8B). Since an increased AMP:ATP ratio is associated with reduced proliferation (Hardie, 2011; Wang et al., 2003) as well as reduced cell viability (Maldonado and Lemasters, 2014), these results confirm the reduced ATP generation in macrophages without GM-CSF stimulation and support the conclusion that GM-CSF is a critical regulator of mitochondrial functions critical to macrophage proliferation and cell survival.

**Fig. 8.**
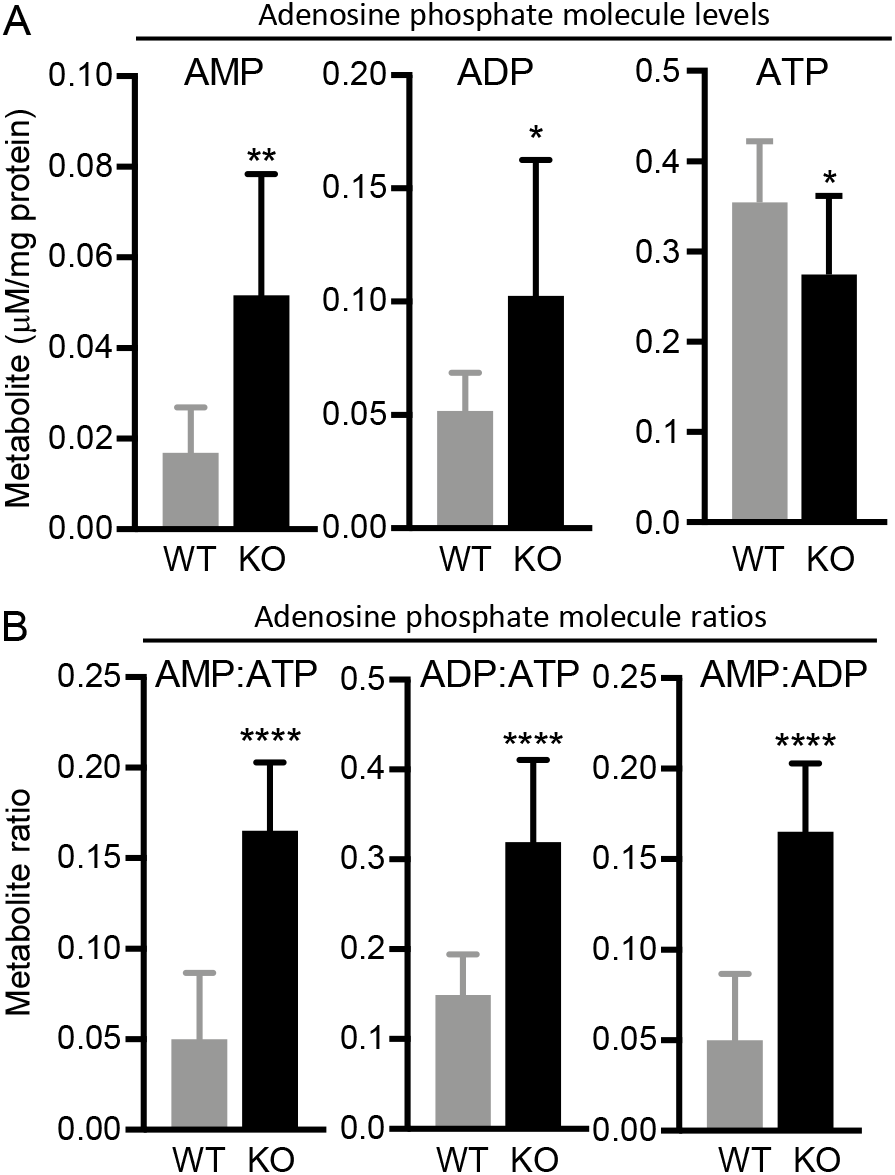
Regulation of energy metabolism by GM-CSF. (A) Adenosine phosphate molecules AMP, ADP and ATP and (B) ratios of adenosine phosphate molecules. Abbreviations: AMP, adenosine monophosphate; ADP, adenosine diphosphate; ATP, and adenosine triphosphate; ns - not significant, *P<0.05, **P<0.01, ****P<0.0001.

## 4. Discussion

In this report, we show that disruption of GM-CSF signaling resulted in multiple mitochondrial abnormalities including undetectable fatty acid metabolism, reduced TCA cycle activity, reduced oxidative phosphorylation, reduced production of ROS and ATP, and a reduced mitochondrial membrane potential and, paradoxically, increased mitochondrial mass – suggesting mitophagy is impaired. Further, loss of GM-CSF signaling was associated with reduced amino acid biosynthesis, glutathione formation, and glycolysis, as well as widespread changes in the expression of genes involved in metabolic pathways including TCA cycle, oxidative phosphorylation, glycolysis, and the pentose phosphate pathway. These results indicate GM-CSF exerts pleiotropic effects on macrophages through critical regulation of mitochondrial turnover, function, and integrity (Figure 9).

**Fig. 9.**
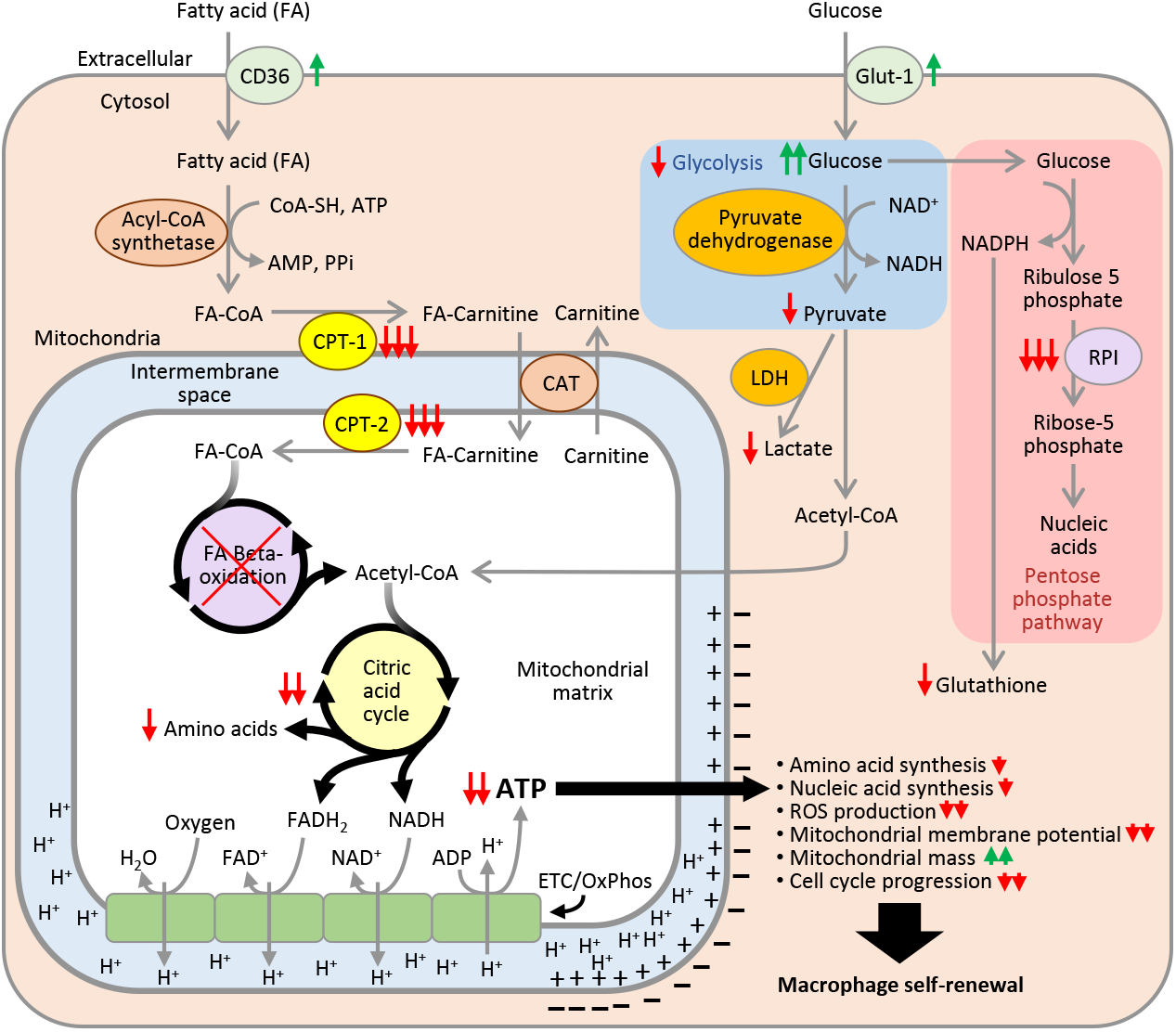
GM-CSF is important for mitochondrial turnover, integrity and functions as well as various cellular cytosolic pathways which are relevant to alveolar macrophage self-renewal. Disruption of GM-CSF signaling in macrophages results in decreased glycolysis, TCA cycle activity, oxidative phosphorylation, ATP production, amino acid and nucleotide precursor synthesis. In addition, fatty acid oxidation is completely abolished due to the downregulation of the mitochondrial enzymes (CPT-1 and CPT-2) responsible for transporting long-chain fatty acids from cytosol to mitochondria for oxidation. Abbreviations: CD36, cluster of differentiation 36; RPI, ribose 5-phosphate isomerase; ROS, reactive oxygen species; CoA-SH, coenzyme A; ATP, adenosine triphosphate; AMP, adenosine monophosphate; PPi, inorganic pyrophosphate; FA-CoA, fatty-acyl CoA; FA-carnitine, Fatty acyl carnitine; CPT-1, Carnitine palmitoyl transferase 1; CPT-2, Carnitine palmitoyl transferase 2; CAT, Carnitine acyl CoA carnitine translocase; H_2_0, water; FAD+, Flavin adenine dinucleotide; FADH2, Flavin adenine dinucleotide reduced; NAD+, nicotinamide adenine dinucleotide; NADH, nicotinamide adenine dinucleotide reduced; H+, hydrogen ion; ETC/OxPhos, electron transport chain/oxidative phosphorylation; LDH, Lactate dehydrogenase; Glut-1, glucose transporter-1. Green vertical arrows indicate increases in pathway/gene expression and/or metabolites and red vertical arrows indicate decreases. Red X indicate that the FA-β-oxidation pathway is undetectable.

The observation that metabolism was severely reduced in macrophages in the absence of GM-CSF signaling has important implications for cellular proliferation but also for viability and the numerous housekeeping and host defense functions for which macrophages are responsible. While glycolysis was moderately reduced (by about half), fatty acid catabolism was abolished, suggesting that, in macrophages, GM-CSF may have a more important role in regulating the catabolism of fatty acids than of glucose. This interpretation is supported by the observation that normal macrophages had a low glycolytic reserve and GM-CSF signaling-deficient macrophages had very little ATP production from palmitate utilization and no detectible spare capacity through fatty acid catabolism. Further, for AMs, this interpretation is consistent with the low concentration of glucose normally present in lung epithelial lining fluid (Baker et al., 2007; Valeyre et al., 1991) and also with their constitutive responsibility for clearing fatty acid-containing phospholipids from the alveolar surface. Lastly, it suggests the possibility that AMs may normally derive more of their energy from the catabolism of fatty acid-containing surfactant phospholipids than from glucose. One potential explanation for the inability of GM-CSF signaling-deprived macrophages to metabolize fatty acids is the lack of expression of *CPT1a, CPT2*, and *FABP1* (Suzuki et al., 2014), which facilitate translocation of fatty acids into mitochondria where they are metabolized via β-oxidation (Jones and Bennett, 2017). Another is the impairment of mitochondrial integrity and function that occurs in macrophages devoid of GM-CSF stimulation. It is also possible that the reduced availability of ATP may contribute to the metabolic defects through frustration of rate-limiting ATP-dependent reactions. The increased levels of Glut1, Glut3, and CD36 in *CsfZrb*^*−/−*^ macrophages suggests a possible (although admittedly speculative) compensatory physiological cellular response aimed at increasing substrate availability in an effort to increase glycolysis and fatty acid oxidation.

The observed reduction in amino acids derived from glycolysis (pyruvate, alanine, valine, leucine, glycine), TCA cycle (aspartate, glutamate, methionine, threonine and isoleucine), and the pentose pathway (histidine) was supported by marked alterations in the transcriptome for genes related to these pathways. Reduced expression of the non-oxidative (anabolic) phase of the pentose pathway suggests that production of ribose-5-phosphate may also be impaired in GM-CSF signaling deficient macrophages. The striking absence of phenylalanine and histidine and reductions in isoleucine, leucine, threonine and valine are important since they comprise the essential amino acids. In addition, reduced levels of glycolytic and TCA cycle intermediates/derivatives that contribute to histone epigenetic modifications including methylation (glycine and betaine), acetylation (acetate) and phosphorylation (AMP/ATP and ADP/ATP ratios) suggest a role for GM-CSF in regulating chromatin modifications during macrophage proliferation. Together, these changes suggest that GM-CSF normally stimulates the synthesis of multiple cellular building blocks required for ATP, protein and DNA synthesis that are required for cellular proliferation.

The observation that the genes involved in oxidative phosphorylation that were downregulated are located in the mitochondrial DNA while those that were upregulated are encoded in the non-mitochondrial DNA was particularly interesting and supports a vital role for GM-CSF in mitochondrial function. This concept is further supported by the observation that, in the absence of GM-CSF stimulation, mitochondrial membrane potential, oxidative phosphorylation, and ATP production are all severely reduced and mitochondrial ultrastructure is disrupted. The marked increase in mitochondrial mass is interesting although the mechanism responsible is unclear.

Peroxisome proliferator-activated receptor γ (PPARγ) and PPARγ coactivator 1α (PGC1 α) play crucial roles in the coordinated regulation of energy metabolism through their effects on fatty acid oxidation, glucose transport, mitochondrial and peroxisomal remodeling, mitochondrial biogenesis, and oxidative phosphorylation (Austin and St-Pierre, 2012). Our current observation of reduced PGC1 α expression and decreased mitochondrial membrane potential in the absence of GM-CSF signaling points towards defective mitochondrial clearance (mitophagy). Our previous report in *Csf2rb−/−* AMs showed decreased *Park2* expression but with no change in the *Pink*1 levels further supporting our theory that GM-CSF signaling is required for mitophagy, essential to maintain mitochondrial quality control. Accumulation of dysfunctional mitochondria has significant implications in cellular proliferation and long-term maintenance. In HSCs, accumulation of dysfunctional mitochondria has been shown to confer divisional memory necessary for their regenerative potential (Hinge et al., 2020). Decreased expression of PGC1a/PPARγ in pulmonary artery smooth muscle cells led to derangements in mitochondrial structure and function (Yeligar et al., 2018).

The observation that GM-CSF signaling is critical to the proliferation of macrophages is further supported by the findings in *Csf2rb*^*−/−*^ mice showing reduced glutathione and cellular ROS levels, and greater percentage of cells arrested in the G0/G1 phase of the cell cycle and thus compromised cell division. Multiple lines of evidence from this study suggest that decreased ATP generation in *Csf2rb*^*−/−*^ macrophages leads to increases in AMP-ADP/ATP ratios. The increased ratios of adenosine nucleoside phosphates is known to stimulate AMP-activated protein kinase (AMPK), an energy sensor critical to the maintenance of cellular energy homeostasis. In addition, AMPK has been reported to regulate key cellular functions including regulation of mitochondrial biogenesis, mitophagy, cell polarity, cell growth and proliferation which are all critical for self-renewal of macrophages (Canto et al., 2010; Davies et al., 1995; Fogarty and Hardie, 2009; Foretz et al., 1998; Hardie, 2011; Jager et al., 2007; Leclerc et al., 1998; Lin et al., 2010; Metallo et al., 2009; Zhang et al., 2006; Zheng and Cantley, 2007; Zong et al., 2002). Another possible explanation that supports our findings is that mitochondrial depolarization and decreased ATP production specifically blocks the G1-to-S cell cycle progression due to decreased levels of cyclin E (Finkel and Hwang, 2009; Mitra et al., 2009). Similar finding in endothelial progenitor cells suggest that GM-CSF stimulation results in increased levels of cyclin D1 and E facilitating cell cycle progression from G1-S phase thereby accelerating proliferation and viability (Qiu et al., 2014). Furthermore, in the absence of GM-CSF signaling reduced amino acid and nucleotide precursors could result in decreased biomass production required for cell division.

The limitations of this study include the fact that we did not determine the relative importance and contributions of each of the various metabolic pathways to cellular proliferation. Given the pleiotropic effects of GM-CSF, we did not determine the inter-gene network regulating the metabolic and mitochondrial alterations observed during GM-CSF blockade.

## 5 Conclusion

We conclude GM-CSF exerts it pleotropic effects on macrophages in part through a critical role in maintaining mitochondria. Results showed disruption of GM-CSF signaling impaired mitochondrial turnover, functions, and integrity, cellular metabolism (tricarboxylic acid cycle, glycolysis, and pentose phosphate pathways), energy production, amino acid and nucleotide synthesis, ROS production, the cell redox-state, and was required for macrophages to exit G_0_/G_1_ and progress through cell division. These protean manifestations are expected to profound effects on macrophage self-renewal.

## Acknowledgements

This work was supported by the CCHMC Foundation Trustee Grant (to PA). We thank D. Black for the assistance in *Csf2rb−/−* mouse colony breeding and maintenance. We would like to acknowledge and thank the assistance from the Research Flow Cytometry Core, Veterinary services and Confocal Imaging Core at Cincinnati Children’s Hospital Medical Center.

---

**Fig. S1.**
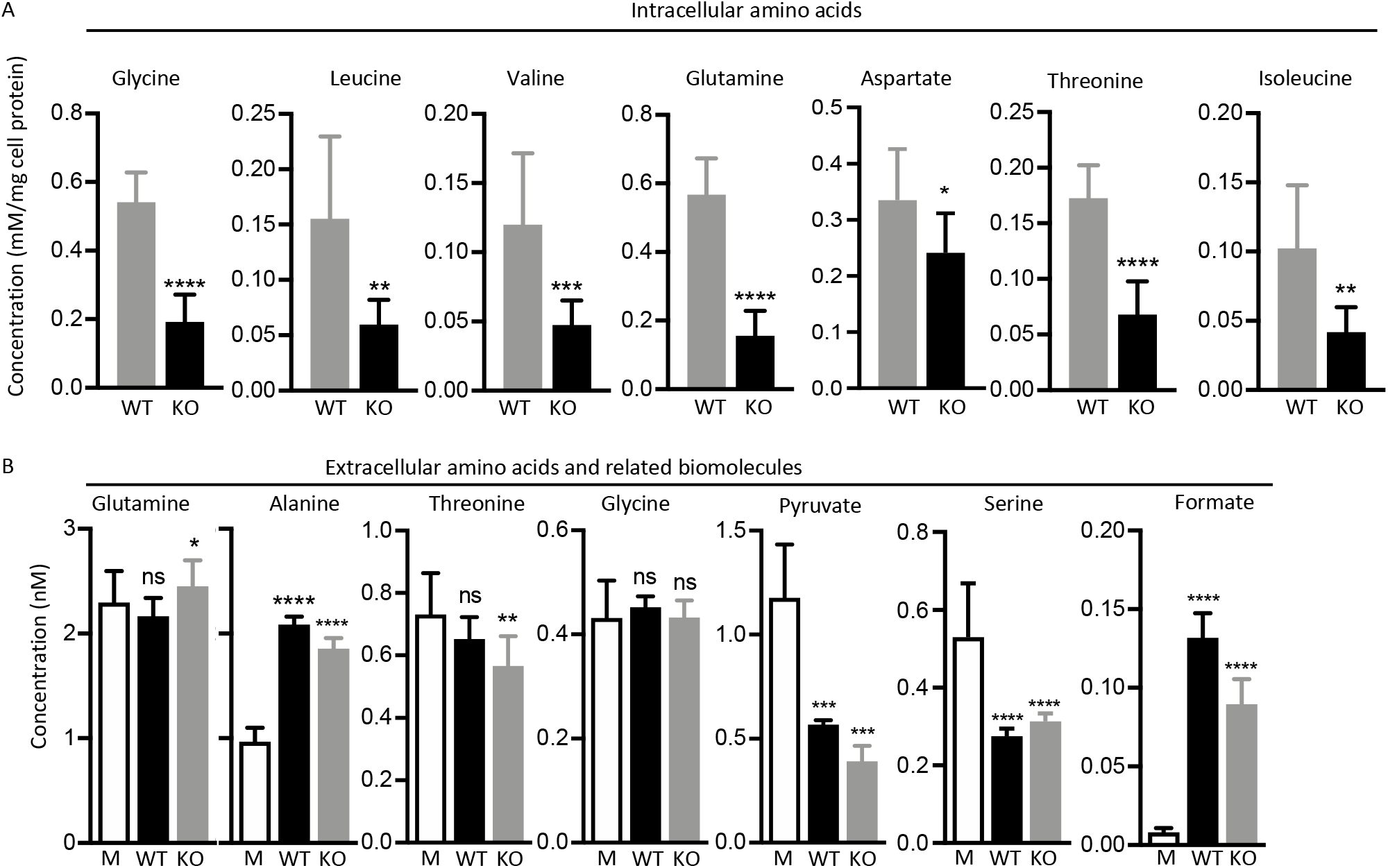
Quantitative NMR metabolite profiles of intracellular and extracellular amino acids and its derivatives in WT and *CsfZrb*^*−/−*^ macrophages. (A) Concentrations of intracellular amino acids glycine, leucine, valine, glutamine, aspartate, threonine and isoleucine. (B) Concentrations of extracellular amino acids and derivatives glutamine, alanine, threonine, glycine, pyruvate, serine and formate, ns, not significant, *P<0.05, **P<0.01, ***P<0.001, ****P<0.0001.

## Notes

### Competing Interest Statement

The authors have declared no competing interest.

## References

Arumugam, P., Suzuki, T., Shima, K., McCarthy, C., Sallese, A., Wessendarp, M., Ma, Y., Meyer, J., Black, D., Chalk, C., Carey, B., Lachmann, N., Moritz, T., Trapnell, B.C., 2019. Long-Term Safety and Efficacy of Gene-Pulmonary Macrophage Transplantation Therapy of PAP in Csf2ra(−/−) Mice. Mol Ther 27, 15971611.

Austin, S., St-Pierre, J., 2012. PGC1alpha and mitochondrial metabolism--emerging concepts and relevance in ageing and neurodegenerative disorders. J Cell Sci 125, 4963–4971.

Aziz, A., Soucie, E., Sarrazin, S., Sieweke, M.H., 2009. MafB/c-Maf deficiency enables self-renewal of differentiated functional macrophages. Science 326, 867–871.

Baker, E.H., Clark, N., Brennan, A.L., Fisher, D.A., Gyi, K.M., Hodson, M.E., Philips, B.J., Baines, D.L., Wood, D.M., 2007. Hyperglycemia and cystic fibrosis alter respiratory fluid glucose concentrations estimated by breath condensate analysis. J Appl Physiol (1985) 102, 1969–1975.

Basu, S., Broxmeyer, H.E., Hangoc, G., 2013. Peroxisome proliferator-activated-gamma coactivator-1alpha-mediated mitochondrial biogenesis is important for hematopoietic recovery in response to stress. Stem Cells Dev 22, 1678–1692.

Berclaz, P.Y., Carey, B., Fillipi, M.D., Wernke-Dollries, K., Geraci, N., Cush, S., Richardson, T., Kitzmiller, J., O’Connor, M., Hermoyian, C., Korfhagen, T., Whitsett, J.A., Trapnell, B.C., 2007. GM-CSF regulates a PU.1-dependent transcriptional program determining the pulmonary response to LPS. Am J Respir Cell Mol Biol 36, 114–121.

Berclaz, P.Y., Shibata, Y., Whitsett, J.A., Trapnell, B.C., 2002a. GM-CSF, via PU.1, regulates alveolar macrophage Fcgamma R-mediated phagocytosis and the IL-18/IFN-gamma -mediated molecular connection between innate and adaptive immunity in the lung. Blood 100, 4193–4200.

Berclaz, P.Y., Zsengeller, Z., Shibata, Y., Otake, K., Strasbaugh, S., Whitsett, J.A., Trapnell, B.C., 2002b. Endocytic internalization of adenovirus, nonspecific phagocytosis, and cytoskeletal organization are coordinately regulated in alveolar macrophages by GM-CSF and PU.1. J Immunol 169, 6332–6342.

Bonfield, T.L., Raychaudhuri, B., Malur, A., Abraham, S., Trapnell, B.C., Kavuru, M.S., Thomassen, M.J., 2003. PU.1 regulation of human alveolar macrophage differentiation requires granulocyte-macrophage colony-stimulating factor. Am J Physiol Lung Cell Mol Physiol 285, L1132–1136.

Canto, C., Jiang, L.Q., Deshmukh, A.S., Mataki, C., Coste, A., Lagouge, M., Zierath, J.R., Auwerx, J., 2010. Interdependence of AMPK and SIRT1 for metabolic adaptation to fasting and exercise in skeletal muscle. Cell Metab 11, 213–219.

Carey, B., Staudt, M.K., Bonaminio, D., van der Loo, J.C., Trapnell, B.C., 2007. PU.1 redirects adenovirus to lysosomes in alveolar macrophages, uncoupling internalization from infection. J Immunol 178, 2440–2447.

Chong, J., Wishart, D.S., Xia, J., 2019. Using MetaboAnalyst 4.0 for Comprehensive and Integrative Metabolomics Data Analysis. Current Protocols in Bioinformatics 68, e86.

Cui, Q., Lewis, I.A., Hegeman, A.D., Anderson, M.E., Li, J., Schulte, C.F., Westler, W.M., Eghbalnia, H.R., Sussman, M.R., Markley, J.L., 2008. Metabolite identification via the Madison Metabolomics Consortium Database. Nature biotechnology 26, 162–164.

Cunningham, J.T., Rodgers, J.T., Arlow, D.H., Vazquez, F., Mootha, V.K., Puigserver, P., 2007. mTOR controls mitochondrial oxidative function through a YY1-PGC-1alpha transcriptional complex. Nature 450, 736–740.

Davies, S.P., Helps, N.R., Cohen, P.T., Hardie, D.G., 1995. 5’-AMP inhibits dephosphorylation, as well as promoting phosphorylation, of the AMP-activated protein kinase. Studies using bacterially expressed human protein phosphatase-2C alpha and native bovine protein phosphatase-2AC. FEBS Lett 377, 421-425.

de Almeida, M.J., Luchsinger, L.L., Corrigan, D.J., Williams, L.J., Snoeck, H.W., 2017. Dye-Independent Methods Reveal Elevated Mitochondrial Mass in Hematopoietic Stem Cells. Cell Stem Cell 21, 725–729 e724.

Dranoff, G., Crawford, A.D., Sadelain, M., Ream, B., Rashid, A., Bronson, R.T., Dickersin, G.R., Bachurski, C.J., Mark, E.L., Whitsett, J.A., et al., 1994. Involvement of granulocyte-macrophage colony-stimulating factor in pulmonary homeostasis. Science 264, 713–716.

Finkel, T., Hwang, P.M., 2009. The Krebs cycle meets the cell cycle: mitochondria and the G1-S transition. Proc Natl Acad Sci U S A 106, 11825–11826.

Fletcher, M., Ramirez, M.E., Sierra, R.A., Raber, P., Thevenot, P., Al-Khami, A.A., Sanchez-Pino, D., Hernandez, C., Wyczechowska, D.D., Ochoa, A.C., Rodriguez, P.C., 2015. I-Arginine depletion blunts antitumor T-cell responses by inducing myeloid-derived suppressor cells. Cancer Res 75, 275–283.

Fogarty, S., Hardie, D.G., 2009. C-terminal phosphorylation of LKB1 is not required for regulation of AMP-activated protein kinase, BRSK1, BRSK2, or cell cycle arrest. J Biol Chem 284, 77–84.

Foretz, M., Carling, D., Guichard, C., Ferre, P., Foufelle, F., 1998. AMP-activated protein kinase inhibits the glucose-activated expression of fatty acid synthase gene in rat hepatocytes. J Biol Chem 273, 14767–14771.

Ginhoux, F., Schultze, J.L., Murray, P.J., Ochando, J., Biswas, S.K., 2016. New insights into the multidimensional concept of macrophage ontogeny, activation and function. Nat Immunol 17, 34–40.

Gonzalez-Juarrero, M., Hattle, J.M., Izzo, A., Junqueira-Kipnis, A.P., Shim, T.S., Trapnell, B.C., Cooper, A.M., Orme, I.M., 2005. Disruption of granulocyte macrophage-colony stimulating factor production in the lungs severely affects the ability of mice to control Mycobacterium tuberculosis infection. J Leukoc Biol 77, 914–922.

Hardie, D.G., 2011. AMP-activated protein kinase: an energy sensor that regulates all aspects of cell function. Genes Dev 25, 1895–1908.

Hashimoto, D., Chow, A., Noizat, C., Teo, P., Beasley, M.B., Leboeuf, M., Becker, C.D., See, P., Price, J., Lucas, D., Greter, M., Mortha, A., Boyer, S.W., Forsberg, E.C., Tanaka, M., van Rooijen, N., Garcia-Sastre, A., Stanley, E.R., Ginhoux, F., Frenette, P.S., Merad, M., 2013. Tissue-resident macrophages self-maintain locally throughout adult life with minimal contribution from circulating monocytes. Immunity 38, 792-804.

Hinge, A., He, J., Bartram, J., Javier, J., Xu, J., Fjellman, E., Sesaki, H., Li, T., Yu, J., Wunderlich, M., Mulloy, J., Kofron, M., Salomonis, N., Grimes, H.L., Filippi, M.D., 2020. Asymmetrically Segregated Mitochondria Provide Cellular Memory of Hematopoietic Stem Cell Replicative History and Drive HSC Attrition. Cell Stem Cell 26, 420–430 e426.

Huang, S.C., Everts, B., Ivanova, Y., O’Sullivan, D., Nascimento, M., Smith, A.M., Beatty, W., Love-Gregory, L., Lam, W.Y., O’Neill, C.M., Yan, C., Du, H., Abumrad, N.A., Urban, J.F., Jr., Artyomov, M.N., Pearce, E.L., Pearce, E.J., 2014. Cell-intrinsic lysosomal lipolysis is essential for alternative activation of macrophages. Nat Immunol 15, 846–855.

Hui, X., Gu, P., Zhang, J., Nie, T., Pan, Y., Wu, D., Feng, T., Zhong, C., Wang, Y., Lam, K.S., Xu, A., 2015. Adiponectin Enhances Cold-Induced Browning of Subcutaneous Adipose Tissue via Promoting M2 Macrophage Proliferation. Cell Metab 22, 279–290.

Ito, K., Carracedo, A., Weiss, D., Arai, F., Ala, U., Avigan, D.E., Schafer, Z.T., Evans, R.M., Suda, T., Lee, C.H., Pandolfi, P.P., 2012. A PML-PPAR-delta pathway for fatty acid oxidation regulates hematopoietic stem cell maintenance. Nat Med 18, 1350–1358.

Ito, K., Ito, K., 2018. Hematopoietic stem cell fate through metabolic control. Exp Hematol 64, 1–11.

Ito, K., Suda, T., 2014. Metabolic requirements for the maintenance of self-renewing stem cells. Nat Rev Mol Cell Biol 15, 243–256.

Ito, K., Turcotte, R., Cui, J., Zimmerman, S.E., Pinho, S., Mizoguchi, T., Arai, F., Runnels, J.M., Alt, C., Teruya-Feldstein, J., Mar, J.C., Singh, R., Suda, T., Lin, C.P., Frenette, P.S., Ito, K., 2016. Self-renewal of a purified Tie2+ hematopoietic stem cell population relies on mitochondrial clearance. Science 354, 11561160.

Jager, S., Handschin, C., St-Pierre, J., Spiegelman, B.M., 2007. AMP-activated protein kinase (AMPK) action in skeletal muscle via direct phosphorylation of PGC-1alpha. Proc Natl Acad Sci U S A 104, 12017–12022.

Janzen, V., Forkert, R., Fleming, H.E., Saito, Y., Waring, M.T., Dombkowski, D.M., Cheng, T., DePinho, R.A., Sharpless, N.E., Scadden, D.T., 2006. Stem-cell ageing modified by the cyclin-dependent kinase inhibitor p16INK4a. Nature 443, 421–426.

Jones, P.M., Bennett, M.J., 2017. Disorders of mitochondrial fatty acid beta oxidation, Biomarkers in Inborn errors of Metabolism. Science Direct, pp. 87–101.

Katajisto, P., Dohla, J., Chaffer, C.L., Pentinmikko, N., Marjanovic, N., Iqbal, S., Zoncu, R., Chen, W., Weinberg, R.A., Sabatini, D.M., 2015. Stem cells. Asymmetric apportioning of aged mitochondria between daughter cells is required for stemness. Science 348, 340–343.

Keij, J.F., Bell-Prince, C., Steinkamp, J.A., 2000. Staining of mitochondrial membranes with 10-nonyl acridine orange, MitoFluor Green, and MitoTracker Green is affected by mitochondrial membrane potential altering drugs. Cytometry 39, 203–210.

Lang, R.A., Metcalf, D., Cuthbertson, R.A., Lyons, I., Stanley, E., Kelso, A., Kannourakis, G., Williamson, D.J., Klintworth, G.K., Gonda, T.J., et al., 1987. Transgenic mice expressing a hemopoietic growth factor gene (GM-CSF) develop accumulations of macrophages, blindness, and a fatal syndrome of tissue damage. Cell 51, 675–686.

Leclerc, I., Kahn, A., Doiron, B., 1998. The 5′-AMP-activated protein kinase inhibits the transcriptional stimulation by glucose in liver cells, acting through the glucose response complex. FEBS Lett 431, 180–184.

LeVine, A.M., Reed, J.A., Kurak, K.E., Cianciolo, E., Whitsett, J.A., 1999. GM-CSF-deficient mice are susceptible to pulmonary group B streptococcal infection. J Clin Invest 103, 563–569.

Lin, Y.C., Hung, C.M., Tsai, J.C., Lee, J.C., Chen, Y.L., Wei, C.W., Kao, J.Y., Way, T.D., 2010. Hispidulin potently inhibits human glioblastoma multiforme cells through activation of AMP-activated protein kinase (AMPK). J Agric Food Chem 58, 9511–9517.

Liu, X., Cooper, D.E., Cluntun, A.A., Warmoes, M.O., Zhao, S., Reid, M.A., Liu, J., Lund, P.J., Lopes, M., Garcia, B.A., Wellen, K.E., Kirsch, D.G., Locasale, J.W., 2018. Acetate Production from Glucose and Coupling to Mitochondrial Metabolism in Mammals. Cell 175, 502–513 e513.

Luo, Y., Tucker, S.C., Casadevall, A., 2005. Fc- and complement-receptor activation stimulates cell cycle progression of macrophage cells from G1 to S. J Immunol 174, 7226–7233.

Maki, K.A., Burke, L.A., Calik, M.W., Watanabe-Chailland, M., Sweeney, D., Romick-Rosendale, L.E., Green, S.J., Fink, A.M., 2020. Sleep fragmentation increases blood pressure and is associated with alterations in the gut microbiome and fecal metabolome in rats. Physiological Genomics 52, 280–292.

Maldonado, E.N., Lemasters, J.J., 2014. ATP/ADP ratio, the missed connection between mitochondria and the Warburg effect. Mitochondrion 19 Pt A, 78–84.

Manesia, J.K., Xu, Z., Broekaert, D., Boon, R., van Vliet, A., Eelen, G., Vanwelden, T., Stegen, S., Van Gastel, N., Pascual-Montano, A., Fendt, S.M., Carmeliet, G., Carmeliet, P., Khurana, S., Verfaillie, C.M., 2015. Highly proliferative primitive fetal liver hematopoietic stem cells are fueled by oxidative metabolic pathways. Stem Cell Res 15, 715–721.

Matrka, M.C., Watanabe, M., Muraleedharan, R., Lambert, P.F., Lane, A.N., Romick-Rosendale, L.E., Wells, S.I., 2017. Overexpression of the human DEK oncogene reprograms cellular metabolism and promotes glycolysis. PLoS One 12, e0177952.

Menendez-Gutierrez, M.P., Roszer, T., Fuentes, L., Nunez, V., Escolano, A., Redondo, J.M., De Clerck, N., Metzger, D., Valledor, A.F., Ricote, M., 2015. Retinoid X receptors orchestrate osteoclast differentiation and postnatal bone remodeling. J Clin Invest 125, 809–823.

Metallo, C.M., Walther, J.L., Stephanopoulos, G., 2009. Evaluation of 13C isotopic tracers for metabolic flux analysis in mammalian cells. J Biotechnol 144, 167–174.

Mitra, K., Wunder, C., Roysam, B., Lin, G., Lippincott-Schwartz, J., 2009. A hyperfused mitochondrial state achieved at G1-S regulates cyclin E buildup and entry into S phase. Proc Natl Acad Sci U S A 106, 11960–11965.

Nakata, K., Akagawa, K.S., Fukayama, M., Hayashi, Y., Kadokura, M., Tokunaga, T., 1991. Granulocyte-macrophage colony-stimulating factor promotes the proliferation of human alveolar macrophages in vitro. J Immunol 147, 1266–1272.

Pridans, C., Sauter, K.A., Irvine, K.M., Davis, G.M., Lefevre, L., Raper, A., Rojo, R., Nirmal, A.J., Beard, P., Cheeseman, M., Hume, D.A., 2018. Macrophage colony-stimulating factor increases hepatic macrophage content, liver growth, and lipid accumulation in neonatal rats. Am J Physiol Gastrointest Liver Physiol 314, G388–G398.

Qiu, C., Xie, Q., Zhang, D., Chen, Q., Hu, J., Xu, L., 2014. GM-CSF induces cyclin D1 expression and proliferation of endothelial progenitor cells via PI3K and MAPK signaling. Cell Physiol Biochem 33, 784795.

Robb, L., Drinkwater, C.C., Metcalf, D., Li, R., Kontgen, F., Nicola, N.A., Begley, C.G., 1995. Hematopoietic and lung abnormalities in mice with a null mutation of the common beta subunit of the receptors for granulocyte-macrophage colony-stimulating factor and interleukins 3 and 5. Proc Natl Acad Sci U S A 92, 9565–9569.

Sakagami, T., Beck, D., Uchida, K., Suzuki, T., Carey, B.C., Nakata, K., Keller, G., Wood, R.E., Wert, S.E., Ikegami, M., Whitsett, J.A., Luisetti, M., Davies, S., Krischer, J.P., Brody, A., Ryckman, F., Trapnell, B.C., 2010. Patient-derived granulocyte/macrophage colony-stimulating factor autoantibodies reproduce pulmonary alveolar proteinosis in nonhuman primates. Am J Respir Crit Care Med 182, 49–61.

Sallese, A., Suzuki, T., McCarthy, C., Bridges, J., Filuta, A., Arumugam, P., Shima, K., Ma, Y., Wessendarp, M., Black, D., Chalk, C., Carey, B., Trapnell, B.C., 2017. Targeting cholesterol homeostasis in lung diseases. Sci Rep 7, 10211.

Seidel, U., Huebbe, P., Rimbach, G., 2019. Taurine: A Regulator of Cellular Redox Homeostasis and Skeletal Muscle Function. Mol Nutr Food Res 63, e1800569.

Senokuchi, T., Matsumura, T., Sakai, M., Yano, M., Taguchi, T., Matsuo, T., Sonoda, K., Kukidome, D., Imoto, K., Nishikawa, T., Kim-Mitsuyama, S., Takuwa, Y., Araki, E., 2005. Statins suppress oxidized low density lipoprotein-induced macrophage proliferation by inactivation of the small G protein-p38 MAPK pathway. J Biol Chem 280, 6627–6633.

Shibata, Y., Berclaz, P.Y., Chroneos, Z.C., Yoshida, M., Whitsett, J.A., Trapnell, B.C., 2001. GM-CSF regulates alveolar macrophage differentiation and innate immunity in the lung through PU.1. Immunity 15, 557–567.

Sieweke, M.H., Allen, J.E., 2013. Beyond stem cells: self-renewal of differentiated macrophages. Science 342, 1242974.

Soucie, E.L., Weng, Z., Geirsdottir, L., Molawi, K., Maurizio, J., Fenouil, R., Mossadegh-Keller, N., Gimenez, G., VanHille, L., Beniazza, M., Favret, J., Berruyer, C., Perrin, P., Hacohen, N., Andrau, J.C., Ferrier, P., Dubreuil, P., Sidow, A., Sieweke, M.H., 2016. Lineage-specific enhancers activate self-renewal genes in macrophages and embryonic stem cells. Science 351, aad5510.

Suzuki, T., Arumugam, P., Sakagami, T., Lachmann, N., Chalk, C., Sallese, A., Abe, S., Trapnell, C., Carey, B., Moritz, T., Malik, P., Lutzko, C., Wood, R.E., Trapnell, B.C., 2014. Pulmonary macrophage transplantation therapy. Nature 514, 450–454.

Suzuki, T., Maranda, B., Sakagami, T., Catellier, P., Couture, C.Y., Carey, B.C., Chalk, C., Trapnell, B.C., 2011. Hereditary pulmonary alveolar proteinosis caused by recessive CSF2RB mutations. Eur Respir J 37, 201–204.

Suzuki, T., McCarthy, C., Carey, B.C., Borchers, M., Beck, D., Wikenheiser-Brokamp, K.A., Black, D., Chalk, C., Trapnell, B.C., 2020. Increased Pulmonary GM-CSF Causes Alveolar Macrophage Accumulation. Mechanistic Implications for Desquamative Interstitial Pneumonitis. Am J Respir Cell Mol Biol 62, 87–94.

Suzuki, T., Sakagami, T., Rubin, B.K., Nogee, L.M., Wood, R.E., Zimmerman, S.L., Smolarek, T., Dishop, M.K., Wert, S.E., Whitsett, J.A., Grabowski, G., Carey, B.C., Stevens, C., van der Loo, J.C., Trapnell, B.C., 2008. Familial pulmonary alveolar proteinosis caused by mutations in CSF2RA. J Exp Med 205, 2703–2710.

Suzuki, T., Sakagami, T., Young, L.R., Carey, B.C., Wood, R.E., Luisetti, M., Wert, S.E., Rubin, B.K., Kevill, K., Chalk, C., Whitsett, J.A., Stevens, C., Nogee, L.M., Campo, I., Trapnell, B.C., 2010. Hereditary pulmonary alveolar proteinosis: pathogenesis, presentation, diagnosis, and therapy. Am J Respir Crit Care Med 182, 1292–1304.

Takihara, Y., Nakamura-Ishizu, A., Tan, D.Q., Fukuda, M., Matsumura, T., Endoh, M., Arima, Y., Chin, D.W.L., Umemoto, T., Hashimoto, M., Mizuno, H., Suda, T., 2019. High mitochondrial mass is associated with reconstitution capacity and quiescence of hematopoietic stem cells. Blood Adv 3, 2323–2327.

Tanaka, T., Motoi, N., Tsuchihashi, Y., Tazawa, R., Kaneko, C., Nei, T., Yamamoto, T., Hayashi, T., Tagawa, T., Nagayasu, T., Kuribayashi, F., Ariyoshi, K., Nakata, K., Morimoto, K., 2011. Adult-onset hereditary pulmonary alveolar proteinosis caused by a single-base deletion in CSF2RB. J Med Genet 48, 205–209.

Trapnell, B.C., Whitsett, J.A., 2002. Gm-CSF regulates pulmonary surfactant homeostasis and alveolar macrophage-mediated innate host defense. Annu Rev Physiol 64, 775–802.

Uchida, K., Nakata, K., Suzuki, T., Luisetti, M., Watanabe, M., Koch, D.E., Stevens, C.A., Beck, D.C., Denson, L.A., Carey, B.C., Keicho, N., Krischer, J.P., Yamada, Y., Trapnell, B.C., 2009. Granulocyte/macrophage-colony-stimulating factor autoantibodies and myeloid cell immune functions in healthy subjects. Blood 113, 2547–2556.

Uchida, K., Nakata, K., Trapnell, B.C., Terakawa, T., Hamano, E., Mikami, A., Matsushita, I., Seymour, J.F., Oh-Eda, M., Ishige, I., Eishi, Y., Kitamura, T., Yamada, Y., Hanaoka, K., Keicho, N., 2004. High-affinity autoantibodies specifically eliminate granulocyte-macrophage colony-stimulating factor activity in the lungs of patients with idiopathic pulmonary alveolar proteinosis. Blood 103, 1089–1098.

Ulrich, E.L., Akutsu, H., Doreleijers, J.F., Harano, Y., Ioannidis, Y.E., Lin, J., Livny, M., Mading, S., Maziuk, D., Miller, Z., Nakatani, E., Schulte, C.F., Tolmie, D.E., Kent Wenger, R., Yao, H., Markley, J.L., 2008. BioMagResBank. Nucleic acids research 36, D402–408.

Valeyre, D., Soler, P., Basset, G., Loiseau, P., Pre, J., Turbie, P., Battesti, J.P., Georges, R., 1991. Glucose, K+, and albumin concentrations in the alveolar milieu of normal humans and pulmonary sarcoidosis patients. Am Rev Respir Dis 143, 1096–1101.

Wang, W., Yang, X., Lopez de Silanes, I., Carling, D., Gorospe, M., 2003. Increased AMP:ATP ratio and AMP-activated protein kinase activity during cellular senescence linked to reduced HuR function. J Biol Chem 278, 27016–27023.

Wang, Y., Szretter, K.J., Vermi, W., Gilfillan, S., Rossini, C., Cella, M., Barrow, A.D., Diamond, M.S., Colonna, M., 2012. IL-34 is a tissue-restricted ligand of CSF1R required for the development of Langerhans cells and microglia. Nat Immunol 13, 753–760.

Waqas, S.F.H., Hoang, A.C., Lin, Y.T., Ampem, G., Azegrouz, H., Balogh, L., Thuroczy, J., Chen, J.C., Gerling, I.C., Nam, S., Lim, J.S., Martinez-Ibanez, J., Real, J.T., Paschke, S., Quillet, R., Ayachi, S., Simonin, F., Schneider, E.M., Brinkman, J.A., Lamming, D.W., Seroogy, C.M., Roszer, T., 2017. Neuropeptide FF increases M2 activation and self-renewal of adipose tissue macrophages. J Clin Invest 127, 3559.

Willinger, T., Rongvaux, A., Takizawa, H., Yancopoulos, G.D., Valenzuela, D.M., Murphy, A.J., Auerbach, W., Eynon, E.E., Stevens, S., Manz, M.G., Flavell, R.A., 2011. Human IL-3/GM-CSF knock-in mice support human alveolar macrophage development and human immune responses in the lung. Proc Natl Acad Sci U S A 108, 2390–2395.

Wishart, D.S., Tzur, D., Knox, C., Eisner, R., Guo, A.C., Young, N., Cheng, D., Jewell, K., Arndt, D., Sawhney, S., Fung, C., Nikolai, L., Lewis, M., Coutouly, M.A., Forsythe, I., Tang, P., Shrivastava, S., Jeroncic, K., Stothard, P., Amegbey, G., Block, D., Hau, D.D., Wagner, J., Miniaci, J., Clements, M., Gebremedhin, M., Guo, N., Zhang, Y., Duggan, G.E., Macinnis, G.D., Weljie, A.M., Dowlatabadi, R., Bamforth, F., Clive, D., Greiner, R., Li, L., Marrie, T., Sykes, B.D., Vogel, H.J., Querengesser, L., 2007. HMDB: the Human Metabolome Database. Nucleic acids research 35, D521–526.

Yeligar, S.M., Kang, B.Y., Bijli, K.M., Kleinhenz, J.M., Murphy, T.C., Torres, G., San Martin, A., Sutliff, R.L., Hart, C.M., 2018. PPARgamma Regulates Mitochondrial Structure and Function and Human Pulmonary Artery Smooth Muscle Cell Proliferation. Am J Respir Cell Mol Biol 58, 648–657.

Zhang, L., Li, J., Young, L.H., Caplan, M.J., 2006. AMP-activated protein kinase regulates the assembly of epithelial tight junctions. Proc Natl Acad Sci U S A 103, 17272–17277.

Zheng, B., Cantley, L.C., 2007. Regulation of epithelial tight junction assembly and disassembly by AMP-activated protein kinase. Proc Natl Acad Sci U S A 104, 819–822.

Zong, H., Ren, J.M., Young, L.H., Pypaert, M., Mu, J., Birnbaum, M.J., Shulman, G.I., 2002. AMP kinase is required for mitochondrial biogenesis in skeletal muscle in response to chronic energy deprivation. Proc Natl Acad Sci U S A 99, 15983–15987.

